# State-dependent emergence of coupling between oscillatory and aperiodic neural activity

**DOI:** 10.64898/2026.10.29.762761

**Authors:** Yuhan Lu, Zhuoran Li, Hangze Mao, Xinghui He, Dingyang Liu, Junxi Chen, Louis Tao, Zhuo-Cheng Xiao, Xing Tian

**Affiliations:** Division of Arts and Sciences, New York University Shanghai, Shanghai 200122, China; Shanghai Key Laboratory of Brain Functional Genomics (Ministry of Education), School of Psychology and Cognitive Science, East China Normal University, Shanghai 200062, China; NYU-ECNU Institute of Brain and Cognitive Science, New York University Shanghai, Shanghai 200062, China; Center for Quantitative Biology, Academy for Advanced Interdisciplinary Studies, Peking University, Beijing, 100871, China; School of Medical Technology and Information Engineering, Zhejiang Chinese Medical University, Hangzhou, 310053, China; Department of Neurosurgery, Xiangya Hospital, Central South University, Changsha, China; Department of Neurosurgery, Guangzhou Sanjiu Brain Hospital, Guangzhou 510510, China; Center for Bioinformatics, National Laboratory of Protein Engineering and Plant Genetic Engineering, School of Life Sciences, Peking University, Beijing 100871, China; NYU-ECNU Institute of Mathematical Sciences, New York University Shanghai, Shanghai 200062, China

## Abstract

Neural oscillations and aperiodic activity are two quintessential mesoscopic-level features arising from shared circuits, yet what governs their relationship remains unclear. Here we show that coupling between oscillatory frequency and aperiodic exponent is sporadic and state-dependent. Bias-reduced analyses of 3,245 intracranial contacts from 61 awake subjects showed significant coupling in only <13% of contacts per band. A connectome-constrained dynamical model predicted that neither changes of input to local circuit nor removal of long-range connections modulated the coupling; whereas stronger local inhibitory feedback drove covariation between oscillatory frequency and aperiodic exponent. The model’s predictions were supported by three independent experiments: transcranial stimulation shifted each component independently and focal ablation of remote cortical tissues did not change coupling; whereas coupling emerged during seizures that altered local excitatory-inhibitory balance. These results suggest that the relation between periodic and aperiodic neural activity is neither intrinsically independent nor coupled, but determined by local circuit states.

## Main

Periodic oscillations and aperiodic broadband activity are ubiquitous signatures of cortical dynamics across species and electrophysiological recording scales^1-4^. Both have been associated with changes in brain state, cognition, development and diseases^5-11^. In parallel, theoretical studies have sought to explain how periodic and aperiodic activity arise from synaptic kinetics, excitation-inhibition balance, recurrent dynamics, and large-scale network interactions^12-17^. Yet their relationship remains unresolved: some studies report substantial covariation between oscillatory and aperiodic features^18,19^, whereas others find weak or no detectable association^20,21^. One hypothesizes that periodic and aperiodic features may remain approximately decoupled when resting-state brain activity consists mainly of small fluctuations around a steady state^14,22^, while the other hypothesizes that the coupling may emerge as circuit dynamics shift across brain states and dynamical regimes^1,23^. The mixed observations hinder the theoretical investigation and raise one essential question: are aperiodic and periodic features intrinsically independent or coupled during ongoing cortical activity?

Spectral parameterization methods, such as FOOOF (fitting oscillations and one-over-f), have made the question approachable by decomposing a power spectrum into an aperiodic background and oscillatory peaks^10,24^. Yet, because the method estimates both components from the same spectrum through sequential background and peak fitting, errors in one estimate can therefore be compensated by the other, and hence creating artificial covariance between the recovered aperiodic exponent and periodic peak frequency^25-27^. The intuition is analogous to estimating two variables, X and Y, while constraining their sum X + Y: an overestimate of one quantity must be offset by the other; dependence can be introduced by the estimation procedure even when the underlying variables are independent.

Nevertheless, correlations reported from FOOOF-derived features may include genuine biological coupling. We tested the possibility with ground-truth simulations and examined human sEEG using a bias-reduced method, SPLIT (spectral piece-wise linear inference of timescale), which first separates fractal and oscillatory spectra by irregular resampling before fitting their features^28,29^. In the study, we first show that FOOOF can produce spurious correlations between oscillatory peak frequency and aperiodic exponent, whereas SPLIT reveals no population-level coupling in awake sEEG recordings from 3,245 contacts in 61 subjects, with significant coupling confined to a minority of contacts.

Reducing fitting-induced covariance helps assess whether coupling is present, but does not establish how coupling emerges depending on conditions of neural circuits. We therefore considered three circuit-level linking mechanisms for the neural dynamics. First, externally inputs to local circuit with different spectra might couple the output features even when the circuit is unchanged^12,13^. Second, local excitation-inhibition feedback can shape both oscillatory frequency and the aperiodic exponent^15,16^. Third, long-range projections can alter local dynamics through interregional feedback^14^. We incorporated the three hypotheses into a dynamical model constrained by the primate cortical connectome^17^. The model predicted that (i) coupling would be weak when uncorrelated periodic and aperiodic inputs passed through the baseline network, (ii) stronger local inhibitory feedback would generate coupling through damped oscillatory responses to aperiodic input, and (iii) changes in coupling following long-range disconnection would be limited. We tested the three predictions in three separate studies using transcranial electrical stimulation with concurrent sEEG recordings (27 subjects) to manipulate external input spectra, sEEG recordings across seizure states (19 subjects) to examine altered local circuit states^30,31^, and sEEG recordings before and after radiofrequency thermocoagulation (15 subjects) to probe the consequences of focal disconnection. Consistent with the model predictions, stimulation and ablation produced no significant cross-feature effects, whereas coupling in the beta band changed during seizures. These results support a circuit-state-dependent account of coupling between periodic and aperiodic neural activity.

## Results

### Spectral analysis method affects aperiodic-periodic coupling

#### FOOOF reports apparent coupling in the gamma bands

We first asked whether aperiodic and periodic features covary over time in human intracranial recordings. We analyzed sEEG recordings from awake-state 61 subjects (3,245 contacts), applying FOOOF to the power spectrum of each time window (see *Methods*). For each contact, we estimated the aperiodic exponent and the center frequency of the highest-power oscillatory peak within each of five frequency bands (delta-theta, alpha, beta, low gamma and high gamma) in each time window. We then calculated the Spearman correlation between these paired estimates across windows, separately for each contact and band. This yielded at most one correlation coefficient per contact per band. We tested whether coupling differed from zero at the population level.

The estimated population mean correlation was significantly positive in both low gamma (*n* = 2,524 valid contacts, *β* = 0.152, *p* = 1.13 × 10^-62^; linear mixed-effect [lme] model) and high gamma (*n* = 1,795, *β* = 0.081, *p* = 6.16 × 10^-10^, **Fig. 1a4, a5**). Mean correlations did not differ significantly from zero in delta-theta (*n* = 2,538, *β* = -0.009, *p* = 1.00), alpha (*n* = 879, *β* = 0.007, *p* = 1.00) or beta (*n* = 1,523, *β* = 0.022, *p* = 0.38, **Fig. 1a1-a3**). At the channel level, significant correlations, irrespective of sign, were detected in 41% of low-gamma contacts and 15% of high-gamma contacts. The gamma-band pattern persisted across the tested combinations of spectral window length and temporal averaging width (**Fig. 1b**).

**Figure 1.**
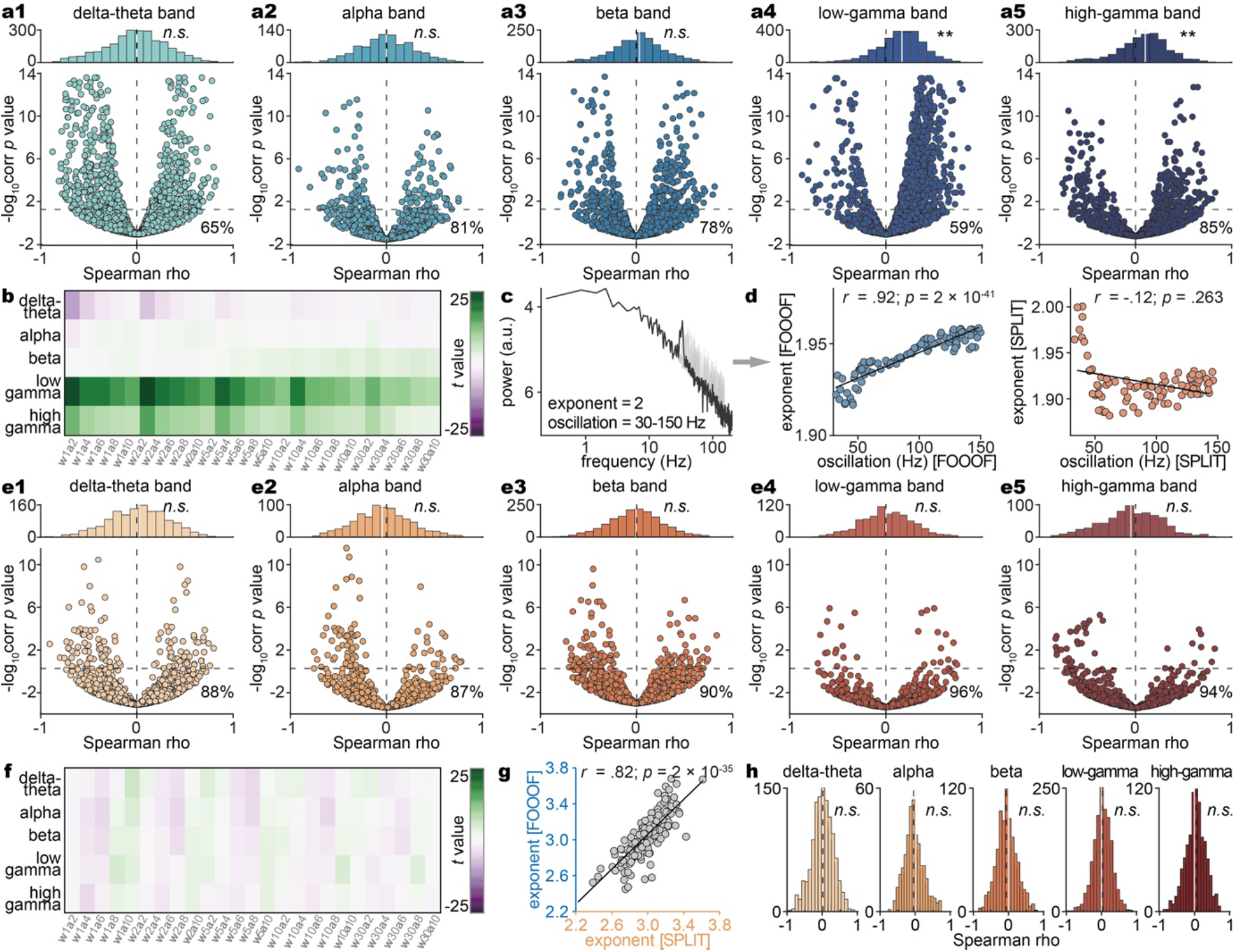
Spectral estimation influences apparent aperiodic-periodic coupling. Awake sEEG recordings from 61 subjects were analysed in **a, b** and **e-h**. Coupling is the within-contact Spearman correlation (ρ) between aperiodic exponent and the centre frequency of the highest-power oscillatory peak across time windows (window length = 5 s, overlap = 50%, number of averaged windows = 6). **a1-a5**, FOOOF estimates in delta-theta (1-8 Hz), alpha (8-13 Hz), beta (13-30 Hz), low gamma (30-80 Hz) and high gamma (80-150 Hz). Histograms show contact-level correlations, with significantly positive population means in low and high gamma. Scatter plots show each correlation and its significance. Percentages indicate contacts without significant correlations. Dashed lines mark ρ = 0 and the significance threshold. **b**, Population-level *t* statistics across 25 analysis settings. Labels *w* and *a* denote spectral-window duration (1, 2, 5, 10 or 30 s) and the number of averaging windows (2, 4, 6, 8 or 10 neighbouring spectra), respectively. **c**, Synthetic spectra combining a fixed Ornstein-Uhlenbeck background, with a high-frequency exponent of 2, and independently generated oscillations at 30-150 Hz. **d**, Exponent and peak-frequency estimates for 100 synthetic signals. FOOOF produced a positive correlation, whereas SPLIT did not detect a significant correlation. Each point represents one signal. **e1-e5**, SPLIT estimates in the recordings, displayed as in **a**. No population mean correlation differed significantly from zero. **f**, Population-level *t* statistics for SPLIT across the same settings. **g**, FOOOF and SPLIT exponent estimates, with one point per recording contact. Lines in **d** and **g** show linear regression fits, with Pearson correlations reported. **h**, Coupling using SPLIT exponents and FOOOF peak frequencies, with no significant population mean correlation. *n*.*s*., not significant. \**p* < 0.05, \*\**p* < 0.005.

Complementary analyses pairing the aperiodic exponent with either the maximum peak power or the highest detected peak frequency also showed apparent coupling (**Fig. S1**). Nevertheless, because FOOOF estimated both parameters from the same spectrum, we next tested whether spectral fitting itself could generate their correlation.

#### Ground-truth simulations reveal spurious coupling in FOOOF estimates

We constructed 100 signals, each combining the same Ornstein-Uhlenbeck (OU) trajectory with an independently generated narrowband oscillation (**Fig. 1c**; see *Methods*). The OU background had a high-frequency exponent of 2, whereas the oscillatory center frequency was drawn randomly from 30 to 150 Hz for each signal. Although only the oscillatory component varied across signals, FOOOF returned a positive exponent-frequency correlation (Pearson *r* = 0.92, *p* = 2×10^-41^, **Fig. 1d**, left). We next applied SPLIT to test whether separating the components before fitting reduced this correlation ^29^. SPLIT first estimates the aperiodic background using irregular-resampling auto-spectral analysis (IRASA), then fits the background and oscillatory peaks separately ^28^. In the same signals, SPLIT yielded no significant correlation between exponent and peak frequency (Pearson *r* = -0.12, *p* = 0.263, **Fig. 1d**, right).

To explain the difference between the two methods, i.e., FOOOF and SPLIT, we quantified how spectral estimation errors propagated to the exponent and peak-frequency estimates (**Fig. S2a, b**). In a representative held-out high-gamma group, FOOOF showed positive coupling, whereas SPLIT showed little coupling (**Fig. S2c**). For FOOOF, second-order predictions derived from calibration simulations tracked the spurious correlations in held-out simulations (Pearson *r* = 0.801, regression slope = 0.693, R^2^ = 0.642, **Fig. S2d**). Spurious coupling was strongest in high gamma (**Fig. S2e**). Subtracting each group’s predicted correlation from its held-out correlation reduced the mean high-gamma Spearman correlation from 0.210 to -0.0160 (**Fig. S2f**). These simulations showed that separating the components before fitting reduced spurious coupling, providing a basis for reassessing the aperiodic-periodic correlations with SPLIT.

#### SPLIT show no consistent coupling across recording contacts

Applying SPLIT to the same intracranial recordings, we found that population mean exponent-frequency correlations did not differ significantly from zero in any band (delta-theta: *n* = 1277, *β* = 0.018, *p* = 1.00; alpha: *n* = 814, *β* = -0.037, *p* = 0.20; beta: *n* = 1616, *β* = -0.009, *p* = 1.00; low-gamma: *n* = 856, *β* = 0.008, *p* = 1.00; high-gamma: *n* = 755, *β* = - 0.045, *p* = 0.15, lme model, FDR corrected, **Fig. 1e**). Significant correlations were detected in fewer than 4% of low-gamma contacts and 6% of high-gamma contacts. This pattern persisted across the tested spectral analysis settings (**Fig. 1f**, in contrast to **Fig. 1b**).

Although FOOOF and SPLIT exponent estimates were strongly correlated across contacts (Pearson *r* = 0.82, *p* = 2 × 10^-35^, **Fig. 1g**), replacing FOOOF exponents with SPLIT exponents while retaining FOOOF peak frequencies yielded no significant population mean correlation in any band (all *p* > 0.114, lme model, FDR corrected, **Fig. 1h**). Our simulation and analytical results suggested that covariance introduced by FOOOF’s interdependent background and peak fitting contributes to the apparent coupling.

So far, our analyses showed that SPLIT detected no significant population mean aperiodic-periodic coupling in awake, intracranial neural recordings, with significant correlations confined to a small minority of contacts. These findings left open, under what conditions, circuit dynamics that shaped these spectral features also made them covary. We therefore used a whole-brain network model to examine how input statistics, local feedback and long-range connectivity affect coupling, and tested the resulting predictions in three separate human experiments.

### A multi-regional network model identifies conditions for periodic-aperiodic coupling

#### Model construction

We developed a general theoretical framework to predict which input and circuit conditions could generate coupling between oscillatory peak frequency and aperiodic exponent.

Because a circuit reshapes the spectrum of its input, we first used a first-order theory that combines input spectral statistics with the circuit’s response at different frequencies to predict output feature correlations (see *Methods*). We then tested this approximation first in simple linear and nonlinear dynamical models (**Figs. S3 and S4**; also see **Supplementary Note**) before applying it to the multi-regional network model constrained by primate cortical connectivity^17^. In the network model, each area contained interacting excitatory and inhibitory populations described by population firing rates, and these local circuits were linked by long-range connections (**Fig. 2a**). For each network condition, we varied the input spectrum across trials and analysed the resulting population activity in the output area. We quantified coupling as the Pearson correlation between output peak frequency and aperiodic exponent across these trials.

**Figure 2.**
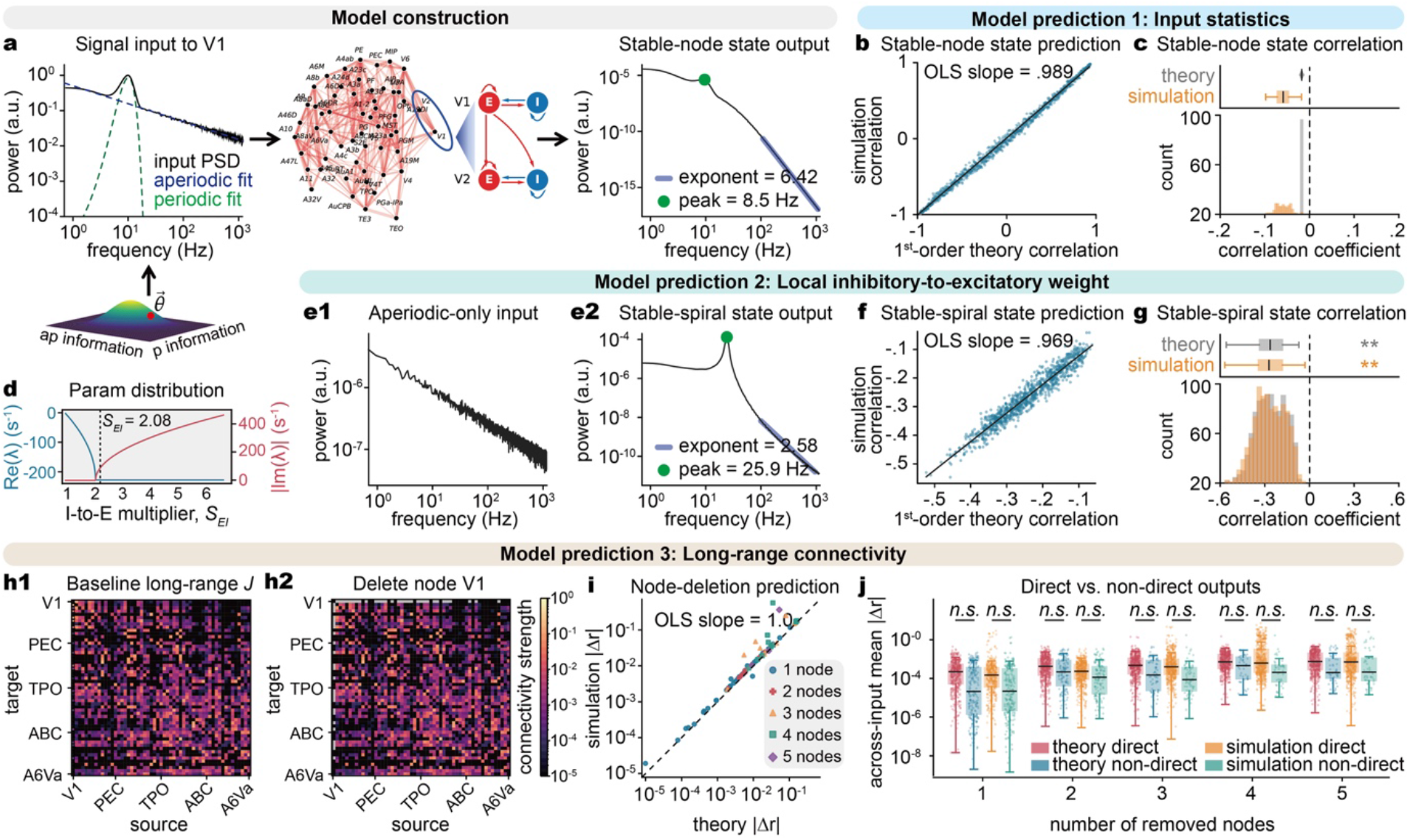
Input and circuit conditions for aperiodic-periodic coupling in a multi-regional network model. Three model predictions concern input statistics, local inhibitory feedback and long-range connectivity. Coupling is the Pearson correlation *r* between output peak frequency and aperiodic exponent across trials. **a**, Oscillatory and aperiodic input delivered to areas in a 55-area excitatory-inhibitory network, with output measured in other areas. Vector θ denotes input spectral parameters. **b**, Predicted and simulated coupling closely matched across 1,000 baseline input conditions. **c**, Weak output correlations across 100 ensembles with uncorrelated input features. No individual simulated correlation was significant. **d**, Real and absolute imaginary parts of local eigenvalues as inhibitory feedback in PG increased, marking a transition from non-oscillatory decay to damped oscillation. **e1-2**, Aperiodic-only input to PG generated an output spectral peak. **f**-**g**, Predicted and simulated coupling closely matched across 1,000 conditions with damped oscillatory responses (**f**), with significant output correlations (**g**). **h1-2**, Long-range connection matrices before and after removing all long-range connections to and from V1 while retaining its local circuit. **i**, Predicted and simulated absolute changes in coupling, |Δ*r*|, closely matched after disconnecting one to five areas. Colours indicate the number of disconnected areas. **j**, Mean |Δ*r*| across input conditions for outputs receiving a direct projection from at least one disconnected area in the intact network and outputs without such projections. Changes were numerically larger for direct outputs, but no difference was significant. Each point in **b** and **f** represents one condition. Lines show ordinary least-squares (OLS) regression fits. Boxes in **c, g** and **j** show medians and interquartile ranges (IQRs), with whiskers spanning values within 1.5 × IQR.

#### Uncorrelated input features yield weak output coupling in the baseline network

We first asked whether the baseline network could generate output coupling when the input features were uncorrelated. With background drive fixed in the non-oscillatory stable-node regime, we delivered inputs containing an oscillatory peak and an aperiodic background to V1 and measured the output in PG. To establish prediction accuracy across different input statistics, we evaluated 1,000 conditions with different correlations between input peak frequency and aperiodic exponent. Predicted and simulated output correlations closely matched (Pearson *r* = 0.998, ordinary least-squares [OLS] slope = 0.989, R^2^ = 0.995; **Fig. 2b**). This agreement extended across all combinations of nine input areas and nine output areas (**Fig. S5**).

We then constructed 100 ensembles in which input peak frequency and aperiodic exponent varied across trials but had numerically zero sample correlation (maximum |r| = 3.82 × 10^-16^). Output correlations were weakly negative, with medians of -0.0174 for the theory and - 0.0587 for simulations (**Fig. 2c**). None of the 100 simulated correlations was individually significant (*p* = 0.44-0.89, two-sided Pearson tests). Nevertheless, the small negative shift was consistent across ensembles, with both theoretical and simulated correlation distributions differing from zero (*p* = 3.90 × 10^-18^ for each, two-sided one-sample Wilcoxon signed-rank tests). Thus, the network produced weak output correlations that were not individually significant. These model findings motivated a transcranial electrical stimulation experiment in humans presented below (**Fig. 3**), to test whether aperiodic stimulation alters oscillatory peak frequency and whether periodic stimulation alters the aperiodic exponent.

**Figure 3.**
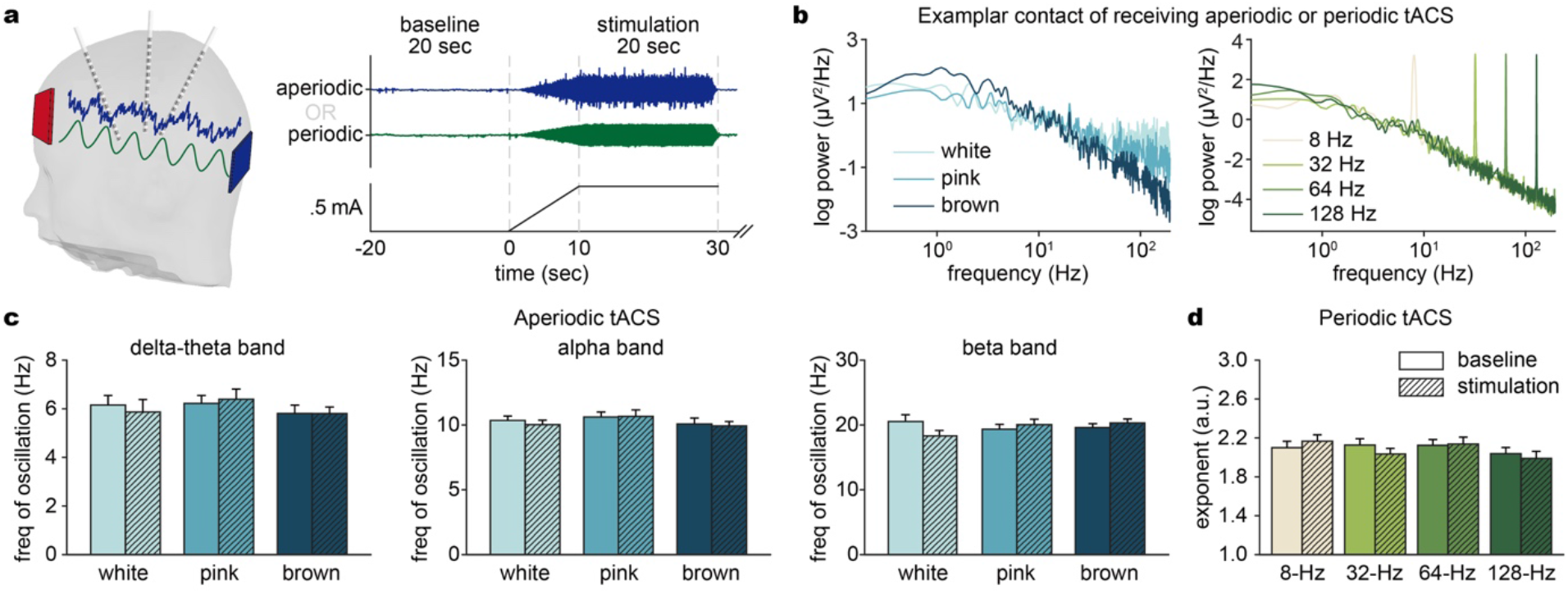
Spectral responses to aperiodic and periodic transcranial stimulation. Transcranial electrical stimulation was combined with simultaneous sEEG recordings in 27 subjects at rest to test whether aperiodic stimulation altered oscillatory peak frequency and periodic stimulation altered aperiodic exponent. **a**, Scalp stimulation, intracranial recording electrodes and stimulation timing. Current increased to 0.5 mA over 10 s and remained at this level for 20 s. Analyses compared the 20-s pre-stimulation baseline with the final 20 s of stimulation. **b**, Example spectra recorded during white-, pink- and brown-noise stimulation (left) and sinusoidal stimulation at 8, 32, 64 and 128 Hz (right). The recorded spectra reflect the applied waveforms in one example contact. **c**, Centre frequency of the highest-power peak in delta-theta (1-8 Hz), alpha (8-13 Hz) and beta (13-30 Hz) before and during aperiodic stimulation. **d**, Aperiodic exponent before and during periodic stimulation, estimated with SPLIT over 0.2-30 Hz. Unhatched and hatched bars indicate baseline and stimulation, respectively. Bars show mean ± SEM across recording contacts. Linear mixed-effects models detected no significant effect of stimulation state on either spectral feature.

#### Local inhibitory feedback can generate coupling from aperiodic input

We next asked whether a change in local circuit dynamics could generate periodic-aperiodic coupling without an oscillatory component in the input. We increased inhibitory feedback onto the excitatory population in area PG while preserving long-range connectivity. Local eigenvalue analysis indicated a transition from non-oscillatory decay to damped oscillations in response to small perturbations at a strength multiplier of approximately 2.08 (**Fig. 2d**). These oscillations decay over time, whereas ongoing aperiodic-only input delivered to V1 generated a peak in the output spectrum at PG (**Fig. 2e1, e2**).

Within each fixed circuit configuration, varying the input aperiodic exponent across trials produced negative correlations between output peak frequency and aperiodic exponent. Across 1,000 parameter conditions, theoretical predictions closely matched simulated correlations (Pearson *r* = 0.964, OLS slope = 0.969, R^2^ = 0.930, **Fig. 2f**). Both theoretical and simulated correlation distributions were significantly negative (*p* = 3.33 × 10^-165^ for each, two-sided one-sample Wilcoxon signed-rank tests, **Fig. 2g**). These results showed that local inhibitory feedback can generate periodic-aperiodic coupling through damped oscillatory responses to aperiodic input. These model findings motivated an analysis of human seizure recordings presented below (**Fig. 4**), to test whether coupling differs across interictal, preictal and ictal periods, given the changes in local excitatory and inhibitory activity associated with seizure transitions ^30,31^.

**Figure 4.**
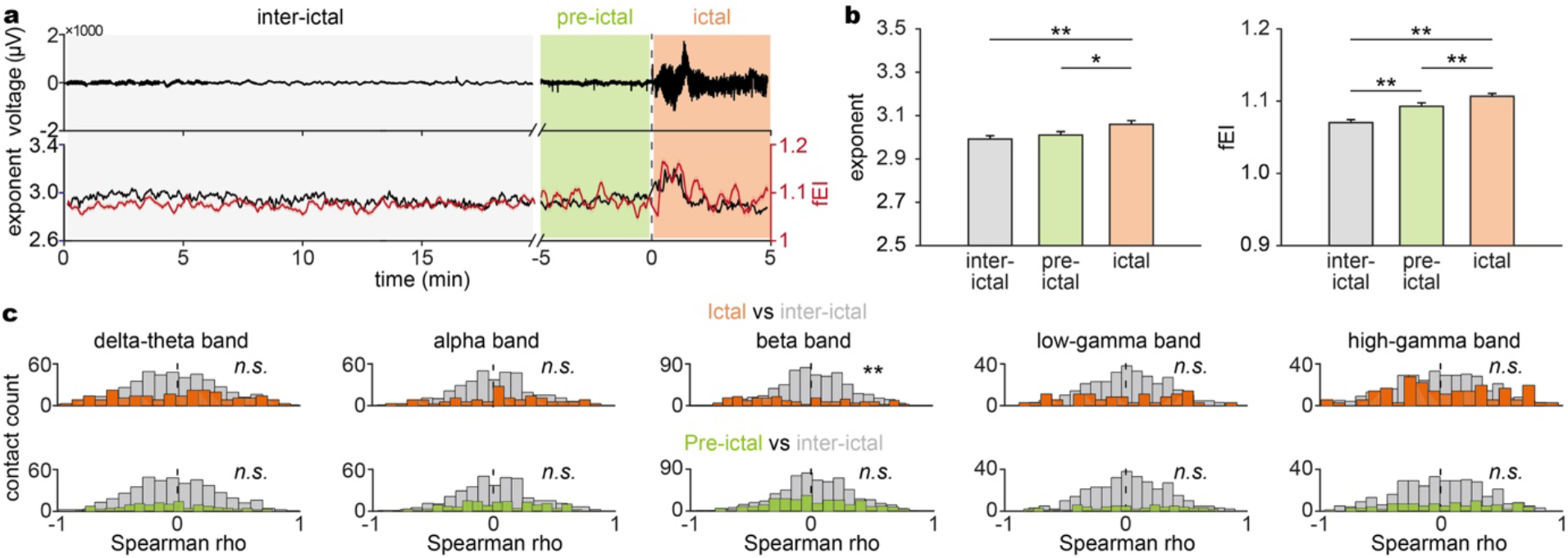
Aperiodic-periodic coupling across seizure states. Coupling and indices of excitation-inhibition balance were examined in stereo-electroencephalography (sEEG) recordings from 19 subjects. **a**, Example sEEG trace and corresponding aperiodic exponent (black) and functional excitation-inhibition ratio (fEI, red). Shading denotes interictal baseline (grey), the 5 min before seizure onset (green) and the ictal period (orange). The dashed line marks seizure onset. **b**, Aperiodic exponent and fEI across states. fEI increased before seizures and further during seizures, whereas the exponent was higher during seizures than during either other period. Bars show mean ± SEM across contacts. **c**, Distributions of coupling during ictal (top) and preictal (bottom) periods, each compared with interictal baseline. Coupling is the within-contact Spearman correlation ρ between exponent and the centre frequency of the highest-power peak across time windows. Only the ictal-interictal beta-band comparison was significant, with coupling shifting towards negative values. Columns show delta-theta (1-8 Hz), alpha (8-13 Hz), beta (13-30 Hz), low gamma (30-80 Hz) and high gamma (80-150 Hz). *n*.*s*., not significant. \**p* < 0.05, \*\**p* < 0.005.

#### Coupling changes after long-range disconnection

We finally asked whether disrupting long-range connections altered periodic-aperiodic coupling in other cortical areas. We disconnected one to five areas by setting their incoming and outgoing long-range weights to zero while retaining their local circuits (**Fig. 2h1, h2**). Using identical input signals, we compared coupling in other cortical areas between the intact and disconnected networks. We quantified the magnitude of these changes as the absolute difference in coupling, |Δr|. Predicted and simulated absolute changes were strongly correlated (Pearson *r* = 0.98, OLS slope = 1.0; **Fig. 2i**).

We compared areas that had received a direct projection from at least one disconnected region (direct outputs) with those that had not (non-direct outputs). Direct outputs showed numerically larger mean absolute changes in coupling than non-direct outputs for all five disconnection sizes in both theory and simulations. However, none of these differences was statistically significant (*p* > 0.170, two-sided sign-flip tests, **Fig. 2j**). These model findings motivated the human radiofrequency thermocoagulation study presented below (**Fig. 5**), which tested whether focal ablation alters coupling at other recording sites and whether these changes differ between sites with and without detectable responses to stimulation at the sites subsequently ablated.

**Figure 5.**
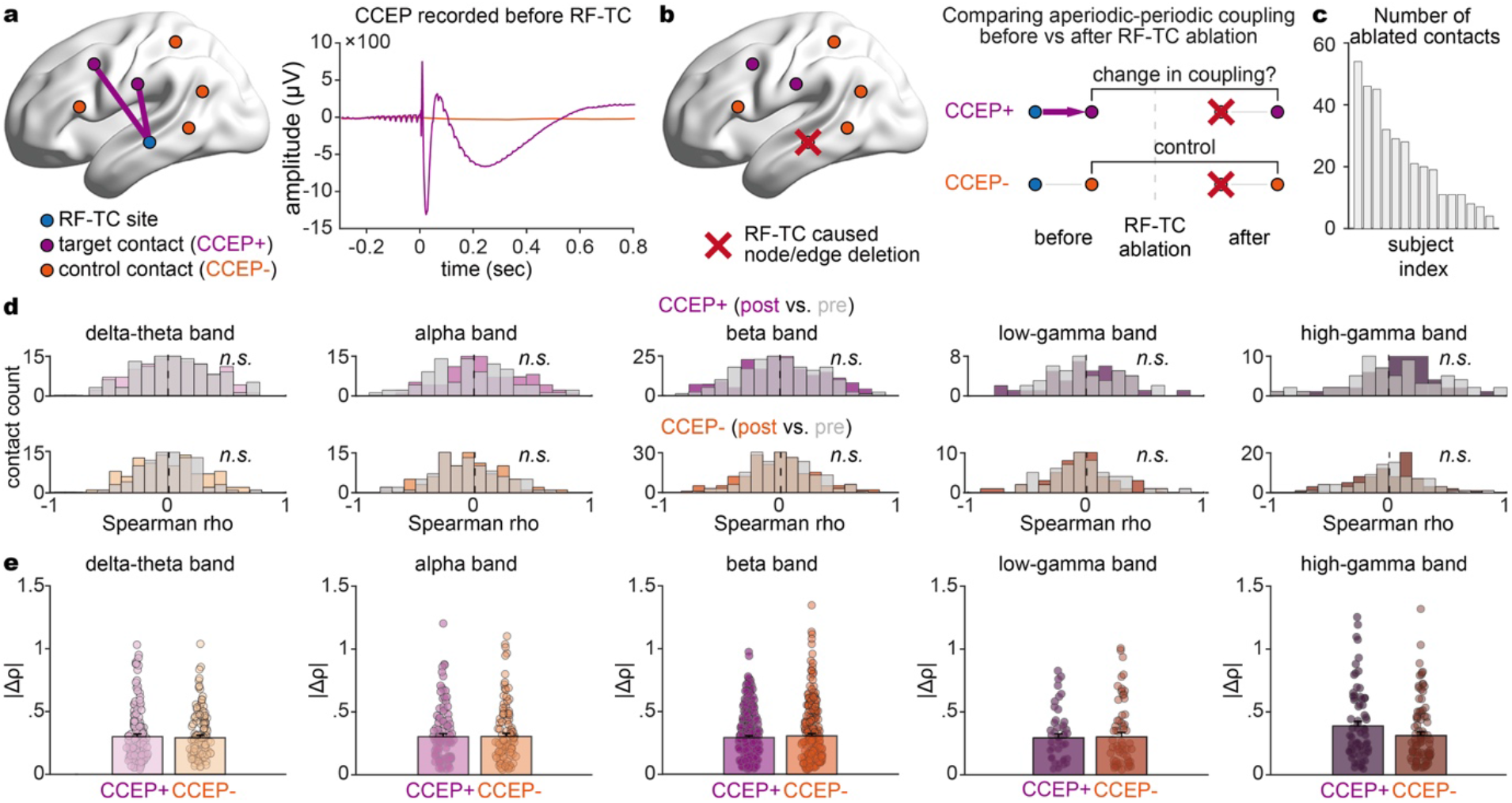
Aperiodic-periodic coupling before and after focal ablation. Stereo-electroencephalography (sEEG) recordings from 15 subjects were used to test whether radiofrequency thermocoagulation (RF-TC) altered coupling at untreated recording sites. **a**, Single-pulse stimulation before RF-TC classified contacts by their cortico-cortical evoked potentials (CCEPs). Contacts responding to at least one subsequently treated site were classified as CCEP+ (purple). Contacts with no detected response to any such site were classified as CCEP-(orange). Blue marks a subsequently treated site, and purple lines indicate detected effective connectivity suggested by CCEPs. Example responses are shown on the right. **b**, Experimental comparison. RF-TC created focal lesions at the sites stimulated in **a** (red crosses). Coupling at untreated CCEP+ and CCEP-contacts was measured in 30-min recordings 2-4 days before and after RF-TC. Coupling was compared before and after RF-TC within each group, and absolute changes were compared between groups. **c**, Number of contacts used for RF-TC in each subject. **d**, Coupling distributions before RF-TC (grey) and after RF-TC in CCEP+ (purple, top) and CCEP-(orange, bottom) contacts. Coupling is the within-contact Spearman correlation ρ between aperiodic exponent and the centre frequency of the highest-power peak across time windows. No significant before-after difference is detected in either group in any band. **e**, Absolute changes in coupling, |Δρ| = |ρ_before_ - ρ_after_|, do not differ significantly between groups. Points represent individual contacts, and bars show mean ± SEM across contacts.

### Empirical tests of the multi-regional model predictions

We examined the three model predictions in three separate human studies of transcranial electrical stimulation (tES), seizure activity and focal ablation.

#### Aperiodic and periodic tES produce no significant cross-component changes

The first study varied the spectrum of the applied current to test whether aperiodic tES altered oscillatory peak frequency and whether periodic tES altered the aperiodic exponent, delivered through frontal and occipital foam electrodes. With simultaneous sEEG recordings in 27 subjects at rest, we compared the 20-s pre-stimulation baseline with the final 20 s of each 30-s stimulation period (**Fig. 3a**).

For aperiodic stimulation, we applied white, pink and brown noise, which differed in spectral slope (**Fig. 3b, left**). We measured the centre frequency of the highest-power peak in each of the delta-theta, alpha and beta bands. Oscillatory peak frequency did not differ significantly between baseline and stimulation (*F*(1, 156) = 0.109, *p* = 0.742, lme model). The stimulation effect did not vary significantly across noise types or frequency bands (all interactions involving stimulation state, *p* > 0.15, **Fig. 3c**). For periodic stimulation, we applied sinusoids at 8, 32, 64 or 128 Hz. Example recorded spectra contained peaks at the stimulation frequencies (**Fig. 3b, right**). Similarly, the aperiodic exponent did not differ significantly between baseline and stimulation (*F*(1, 208) = 1.669, *p* = 0.198, **Fig. 3d**).Thus, neither analysis detected a significant overall change in the other spectral feature during stimulation. These findings were consistent with the weak, nonsignificant output coupling observed in the network model.

#### Aperiodic-periodic coupling changes during seizures

The second model prediction showed that strengthening local inhibitory feedback could generate aperiodic-periodic coupling. Because changes in local excitatory and inhibitory activity have been documented around seizure onset ^30,31^, we examined the prediction by asking whether coupling during preictal and ictal states could be stronger than interictal baseline. In sEEG recordings from 19 subjects, we measured the functional excitation-inhibition ratio (fEI) and aperiodic exponent as proposed indirect indices of excitation-inhibition balance (**Fig. 4a**) ^16,32^. fEI increased before seizure onset and rose further during seizures (*p* = 1.18 × 10^-6^ and 1.10 × 10^-13^, respectively; lme model, FDR corrected; **Fig. 4b**, right). The exponent did not differ significantly between preictal and interictal recordings, but was higher during seizures than during either the interictal or preictal period (*p* = 0.00159 and 0.0406, respectively; **Fig. 4b**, left). These changes in indirect E-I measures suggested altered local circuit states across seizure states.

During the preictal period, coupling did not differ significantly between preictal and interictal recordings in any of the five frequency bands (**Fig. 4c**, bottom). During seizures, the population mean beta-band correlation shifted towards negative values, with the mean Spearman correlation changing from 0.0196 at interictal baseline to -0.1165 during the ictal period (*β* = -0.136, *p* = 0.004, lme model, FDR corrected, **Fig. 4c**, top). Sensitivity test of this beta-band result further confirmed that this ictal-interictal difference was robust to any exclusion of patient subsets (**Table S1**). Therefore, these results showed that aperiodic-periodic coupling significantly changed with seizure state, consistent with the circuit-state dependence suggested by the model.

#### Aperiodic-periodic coupling shows no significant change after focal ablation

The final model prediction showed that areas that had received direct input from disconnected regions had numerically larger changes than the area without such input, although the difference was not statistically significant. Correspondingly, the empirical study compared coupling changes after focal ablation between sites that responded to stimulation of the treated regions and those with no detectable response. Fifteen subjects underwent sEEG-guided radiofrequency thermocoagulation (RF-TC), which creates focal lesions. We grouped the contacts not used for RF-TC according to their responses to stimulation of the subsequently ablated sites. Contacts showing a cortico-cortical evoked potential (CCEP) in response to single-pulse stimulation of at least one such site were classified as CCEP+. Contacts with no detected response to any such site were classified as CCEP-. This classification used recordings obtained before RF-TC (**Fig. 5a**). We then measured coupling before and after RF-TC at these contacts and compared the absolute changes between the two groups (**Fig. 5b,5c**).

Coupling did not differ significantly before and after RF-TC in either CCEP group in any of the five frequency bands (all *p* > 0.211, lme model, FDR corrected, **Fig. 5d**). To assess changes regardless of direction and consistent with the model measures, we also compared the absolute change in coupling, |Δρ|, between CCEP+ and CCEP-contacts. Absolute changes did not differ significantly between the two groups in any band (all *p* > 0.122, lme model, FDR corrected, **Fig. 5e**). Therefore, coupling did not change significantly after RF-TC, and absolute changes did not differ significantly between CCEP+ and CCEP-contacts. This lack of a significant group difference was consistent with the observation in the model after long-range disconnection.

## Discussion

Our findings support a conditional relationship between aperiodic exponent and oscillatory peak frequency. The spectral analysis method, FOOOF, could introduce correlations between these features even when the aperiodic background was unchanged. After reducing this bias, most contacts showed no detectable coupling during spontaneous wakefulness. In our models, strengthening local inhibitory feedback enabled changes in aperiodic input to jointly shift the background slope and peak frequency. The change in beta coupling during seizures is consistent with this sensitivity to altered local excitatory and inhibitory dynamics. These findings suggest that a shared circuit origin does not require the two features to fluctuate together. Rather, their covariance depends on how the circuit’s current state shapes its response to changing input.

Our study identifies a methodological source of apparent coupling between periodic and aperiodic features. Although spectral parameterization helps distinguish oscillatory changes from changes in the background ^10,25^, our simulations show that changing peak frequency can bias the exponent estimated by FOOOF even when the aperiodic background remains unchanged. If peak frequency also varies with behavior, this bias could make the estimated exponent appear behaviorally relevant without a corresponding change in aperiodic activity. The prominence of this artifact in gamma does not imply that it is restricted to gamma. Our controlled simulations show that the affected frequency range depends on knee location. Lower-frequency knees, such as those reported in resting MEG, could therefore shift this bias toward alpha and beta ^33^. These findings motivate checking reported associations between periodic and aperiodic features using simulations matched to the spectral properties of the recordings.

Beyond estimation bias, the physiological question is when shared circuitry makes periodic and aperiodic features vary together. Our mathematical models allow input statistics, local feedback and long-range connectivity to be varied independently, but necessarily simplify the underlying biology. In parallel, our human experiments seek to probe these influences within a more complex biological setting, where their individual contributions cannot be isolated as readily. The models suggest possible mechanisms of spectral coupling, while the experiments help us explore their relevance when multiple processes interact.

Periodic and aperiodic features can arise from shared dynamics. Models of oscillatory relaxation can account for both spectral peaks and broadband scaling^34^, and IRASA analyses have revealed temporal covariance between alpha power and the aperiodic exponent ^18^. Local inhibitory feedback offers a mechanism linking the exponent specifically to peak frequency by organizing nonrhythmic excitation into rhythmic activity^35^, consistent with noise-driven oscillations in stochastic neuronal networks ^36^. In our models, a peak inherited from rhythmic input can vary independently of the background. When local feedback instead produces a peak from aperiodic input, its location depends on both the input spectrum and the circuit’s frequency response. Changes in input spectral shape can then shift the peak and background slope together.

This mechanism provides a hypothesis for the state dependence observed in the seizure recordings. During human mesial temporal low-voltage fast seizures, putative interneurons increase their firing and synchronize with 20-30 Hz activity ^31^. Moreover, activating parvalbumin interneurons in mice can switch from suppressing to promoting seizures within seconds of onset, showing that inhibitory function can change substantially during a seizure ^37^. This supports inhibitory involvement in ictal beta dynamics, although increased interneuron firing does not establish stronger effective inhibitory feedback. We hypothesize that ictal changes in local feedback alter the sensitivity of beta peak frequency to the input spectral variations that also shape the aperiodic background. Changes in this sensitivity could modify how strongly the two features covary, linking the ictal beta effect to the interaction between circuit state and ongoing input.

In conclusion, periodic and aperiodic features can reflect shared circuit dynamics without maintaining a fixed pattern of covariation. This reconciles a common physiological origin with distinct, state-dependent spectral responses. For cognitive functions, it raises the possibility that task-related changes in exponent and peak frequency are different expressions of a shared neural response, whose spectral manifestation depends on the state in which it occurs.

## Methods

### Subjects and study design

This study analyzed intracranial EEG data from patients with medically intractable epilepsy at the Guangdong Sanjiu Brain Hospital (Guangzhou, China) and Xiangya Hospital of Central South University (Changsha, China). The study comprised four datasets from 122 unique patients (see **Table S2** for detailed demographic and clinical information): recordings during wakefulness from 61 patients; simultaneous transcranial electrical stimulation (tES) and sEEG recordings from 27 patients; interictal, preictal, and ictal recordings from 19 patients; and recordings before and after radiofrequency thermocoagulation (RF-TC) from 15 patients. During the wakefulness and RF-TC recordings, patients were awake and responsive and engaged in spontaneous activities, including conversation and phone use. All procedures were approved by the Research Ethics Committee of the two hospitals and the Institutional Review Board of New York University Shanghai.

#### sEEG acquisition and anatomical localization

sEEG was recorded with a 256-channel Neuvo system at 2,000 Hz at Guangdong Sanjiu Brain Hospital and a Natus Quantum system at 2,048 Hz at Xiangya Hospital. Depth electrodes contained 8-16 cylindrical contacts, each 0.8 mm in diameter and 2 mm long, with the gap between contact edges of 1.5 mm (Sinovation, China). Electrode number and placement were determined solely by clinical requirements for presurgical localization of epileptogenic regions. Trajectories were planned using the ROSA stereotactic robotic system and co-registered preoperative MRI and CT images. Postoperative CT images were rigidly co-registered to each patient’s preoperative T1-weighted MRI. Contacts were identified manually on the co-registered images to obtain coordinates in native MRI space. T1-weighted images were segmented and normalized to MNI152 space using CAT12 within SPM12 ^38^. The resulting deformation fields were applied to the contact coordinates. Contacts were anatomically labelled using Brainstorm ^39^.

### sEEG preprocessing

Analyses used custom MATLAB and Python scripts ^40,41^. Continuous recordings were visually screened for epileptiform activity and artifacts by neurologists J.C., X.H., and D.L.. Data were resampled to 1,000 Hz and notch-filtered at 48-52, 98-102, 148-152, and 198-202 Hz. No additional broadband high-pass filtering was applied before spectral estimation.

Signals were then re-referenced by subtracting adjacent contacts along each electrode shaft; the terminal contact did not define an additional bipolar channel. Contacts labelled as white matter or unassigned (N/A) were excluded.

### Time-resolved spectral analysis

#### Spectral parameterization using FOOOF

We evaluated 25 combinations of five Welch window lengths (1, 2, 5, 10 and 30 s) and five temporal averaging widths (2, 4, 6, 8 and 10 neighboring PSD estimates). For each window length, PSD estimates were computed throughout the recording using Welch’s method, with 50% overlap between adjacent windows. Each PSD was smoothed using a centred moving average over the specified neighboring estimates. The averaging width therefore denotes the number of neighboring estimates surrounding the target PSD. The target advanced by one PSD estimate at each step. Each averaged PSD was parameterized with FOOOF over 0.2-200 Hz in knee mode ^10^. Settings were a maximum of three peaks, peak-width limits of 1.5-6 Hz, a minimum peak height of 0.6 and a peak-detection threshold of 2. The aperiodic offset, exponent and knee parameter, together with fitted peak parameters, were retained. In this model, the exponent describes the asymptotic high-frequency decay.

#### Spectral parameterization using SPLIT

##### IRASA decomposition

For comparison with FOOOF, IRASA used the same five window lengths (1, 2, 5, 10 and 30 s), 50% overlap and temporal averaging scheme. IRASA was applied independently to each temporal window to obtain an original spectrum and a fractal spectrum ^28^. It used paired resampling factors h and 1/h (we used default settings h = 1.1, 1.2, …, 1.9). IRASA was computed over 0.1-400 Hz to accommodate the expanded frequency range associated with resampling^25^. Original spectra and fractal spectra were then averaged separately: Each PSD was smoothed using the same moving-average scheme, with 2, 4, 6, 8 or 10 neighboring estimates. This yielded the same 25 combinations of window length and temporal averaging width as the FOOOF analysis. Averaged fractal spectra were used for aperiodic fitting; averaged original and fractal spectra were used together for periodic fitting.

##### Aperiodic fitting

Aperiodic spectra were parameterized using our previously described piecewise-linear fitting procedure ^29^. Spectra were represented in log-log space and interpolated onto 1,000 equally spaced log-frequency points spanning 0.2-200 Hz. Using pwlf, we fitted models with one to four segments, corresponding to zero to three breakpoints. Model order was selected by minimizing

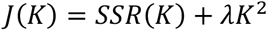

where K is the number of segments, SSR(K) is the residual sum of squares and λ = 0.5. Breakpoints were converted to knee frequencies, and the negative slope of each segment defined its local aperiodic exponent. Fits were visually inspected without manually changing model selection. Coupling analyses were restricted to spectra best fitted by a single-knee model, which enabled direct comparison between FOOOF and SPLIT. Such a model has two segments; we used the negative slope of the second segment as the post-knee aperiodic exponent.

##### Periodic fitting

For each averaged spectrum, the flattened spectrum was calculated as the log10 original PSD minus the log10 fractal PSD. Positive values therefore indicate power above the estimated aperiodic background. Line-noise frequency ranges within ±3 Hz of 50, 100 and 150 Hz were excluded from peak detection and fitting. These bands can contain residual line noise or notch-related artifacts that affect the standard deviation used by the relative peak threshold.

Oscillatory peaks were identified using the Gaussian fit_peaks default procedure, which followed the peak-fitting framework of spectral parameterization^10,24^. Candidate peaks were identified iteratively from the largest residual maximum exceeding both the relative threshold and the minimum peak height. Candidate Gaussian components were subtracted during peak identification, and retained candidates were fitted jointly. The relative peak-detection threshold was 2 residual standard deviations, and the proximity threshold was 2.5 Gaussian standard deviations. Peak-width limits were 1.5-6 Hz. Peaks were fitted separately over 1-30 Hz, with up to three peaks and a minimum height of 0.6, and over 30-80 Hz and 80-150 Hz, with up to one peak per range and a minimum height of 0.8^42^. The reported Gaussian features were centre frequency and amplitude above the aperiodic background in log10 power units. Spectra with no qualifying peaks were classified as having no detectable periodic component under these criteria.

#### Quantification of aperiodic-periodic coupling

For each electrode channel and time window, the aperiodic exponent was paired with the center frequency of each detected peak. Peaks were assigned by center frequency to delta-theta (1-8 Hz), alpha (8-13 Hz), beta (13-30 Hz), low gamma (30-80 Hz) or high gamma (80-150 Hz). When more than one peak occurred within the same frequency band and time window, only the peak with the highest power was retained. Within each channel and band, exponent-frequency pairs were pooled across time. Pairs were excluded if either variable fell below its first quartile minus 1.5 times the interquartile range or above its third quartile plus 1.5 times the interquartile range. When at least 20 pairs remained, coupling was quantified as Spearman’s rank correlation, ρ, between the aperiodic exponent and peak center frequency across time. Each channel therefore contributed at most one coefficient per band for a given recording, state and spectral setting. Complementary analyses also used exponent with the peak with strongest power and with the highest-frequency, shown in **Fig. S1**.

### Ground-truth OU simulation for method validation

#### Simulated signal generation

We varied four input-generating parameters: oscillation centre *c* (Hz), passband width *w* (Hz), nominal oscillatory-to-background power ratio *q* at *c*, and background knee frequency *f*_*k*_ (Hz). For each parameter draw (*c, w, q, f*_*k*_), we generated an Ornstein-Uhlenbeck background *x*(*t*) and an independent narrow-band component *u*(*t*). The discrete background was

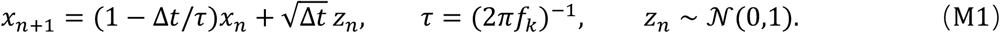

The index *n* denotes time samples, Δ*t* is the sampling interval, and *z*_*n*_ is independent Gaussian white noise. We initialized *x*_0_ = 0. This Euler-Maruyama generator approximates a background with PSD proportional to [τ^−2^ + (2π*f*)^2^]^−1^; the exact discrete spectrum was given in **Supplementary Note S1.1**. To generate the oscillatory component *u*(*t*), we passed independent Gaussian white noise through a second-order Butterworth bandpass filter with passband edges *c* ± *w*/2, applied the filter forwards and backwards, and normalized the result to unit root-mean-square amplitude. The final common input was

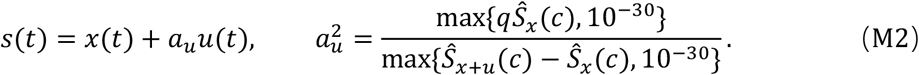

Here Ŝ denotes the Welch PSD and *a*_*u*_ is a time-domain amplitude multiplier from the input parameter *q* to produce the specified oscillatory-to-background power ratio at the centre frequency *c*.

In **Fig. 1c**, signals were 30-s long and sample at 1,000 Hz. We set *f*_*k*_ fixed at 2 Hz and the oscillation centre *c* randomly (uniform distribution) selected from 30 to 150 Hz. We generated 100 composite time series that shared the same OU trajectory and differed in their oscillatory component. In **Fig. S2**, signals were 30-s long and sampled at 1,000 Hz. The parameter range and sampling criteria is in **Supplementary Note S1.2-1.3**. We fitted the simulated spectra with FOOOF and IRASA-SPLIT using the same procedures as for the empirical recordings (see *Time-resolved spectral analysis*). For the analyses in **Fig. S2**, we used calibration simulations to predict correlations between fitted peak frequency and aperiodic exponent, then evaluated these predictions in independent held-out simulations, as described below.

#### First- and second-order prediction

We used perturbation theory to describe how changes in generating parameters alter fitted features. Let **θ** = (*c, w, q, f*_*k*_)^⊤^, let **θ**_0_ be the expansion centre, and let *m*,(**θ**) be the conditional mean estimate of feature *a* ∈ {*P, X*}, where the periodic feature P denotes peak frequency in hertz. The aperiodic feature *X* is the fitted positive exponent. With δ**θ** = **θ** − **θ**_0_,

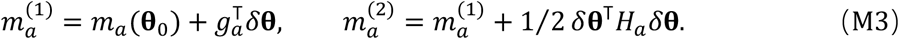

The vector *g*_*a*_ is the gradient and *H*_*a*_ the Hessian of the feature map. The first-order signal covariance is:

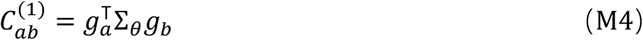

where Σ _θ_ is the input-parameter covariance. Because the input generating parameters in **Fig. S2** are sampled from broad distributions, we estimated these coefficients by count-weighted ridge regression of first- or second-degree response surfaces to calibration means with penalty term (see *Calibration, held-out evaluation and statistical analysis*) Their accuracy was tested on held-out outcomes. The Gaussian second-order covariance identity and its non-Gaussian extension 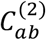 were derived in **Supplementary Note S2**.

To locate spectral contributions, we also linearized each estimator with respect to log-spectral deviations ϵ_*j*_ at frequency *f*_*j*_. Writing 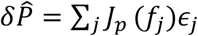 and 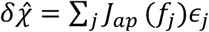 gives the signed frequency-resolved local first-order contributions to *Cov*(*P, A*):

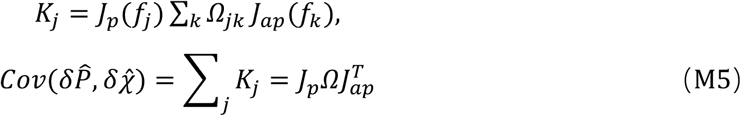

*J*_*p*_, *J*_*ap*_ are local error-transfer weights and Ω is the calibration covariance of the relevant log-spectral deviations. Hence *J*_*p*_(*f*)and *J*_*ap*_(*f*) identify the frequencies at which spectral perturbations shift the estimated peak centre and aperiodic exponent, thereby showing the local direction and strength of error propagation before the covariance structure of the spectral errors is taken into account. **Supplementary Note S3** derived the weights and defines their display normalization.

#### Calibration, held-out evaluation and statistical analysis

We predicted feature correlation by combining covariance of the conditional mean estimates with covariance arising from repeated realizations at fixed input parameters. For the fitted pair 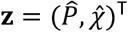 and generating vector **θ** = (*c, w, q, f*_*k*_)^⊤^, the total covariance is

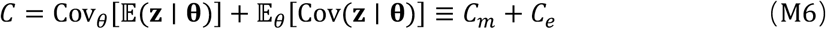

Here *C*_*m*_ describes variation in the conditional mean **m**(**θ**) = E (**z** ∣ **θ**) across input settings. The realization-error covariance *C*_*e*_ describes variation around that mean when the input parameters are fixed. The corresponding Pearson prediction is

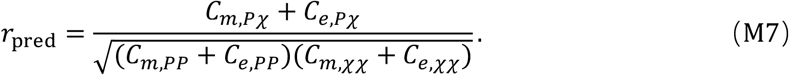

The conditional mean was approximated by the first- or second-order expansion as defined above (M3); the final group comparison used the second-order expansion. Calibration estimated its intercept, gradient *g*_*a*_ and Hessian *H*_*a*_ for each feature *a* ∈ {*P, X*}. For each estimator (FOOOF or IRASA-SPLIT), a realization was valid when that estimator yielded finite estimates of both periodic and aperiodic features. For calibration input parameter setting *i*, we averaged its *n*_=_ valid feature pairs obtained from its realizations:

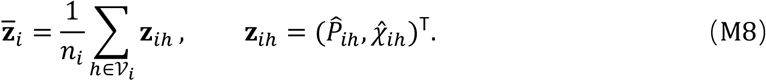

Here *h* indexes realizations and *V*_*i*_ is the set of valid realizations at input parameter setting *i*. We standardized each input parameter using its calibration mean and sample standard deviation and fitted all parameters using count-weighted ridge regression. See **Supplementary Notes S2.2** for more details. Equations (M3) and (M4), together with the regression procedure give the prediction of *C*_*m*_ from calibration trials.

We then estimated realization error from variation within each calibration setting. Subtracting the setting-specific mean from each valid fitted pair produced the residual 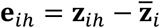. The remaining complete two-dimensional residuals formed an empirical error distribution. Its covariance, used by uniform residual resampling, was

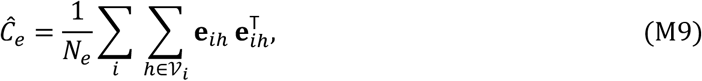

Here *N*_*e*_ is the number of pooled residual pairs. Calibration settings with at least 2 valid realizations were included in sums.

To match the statistical analysis of the experimental data, we used Spearman rank correlation for selected comparisons. Whereas Pearson correlation can be calculated directly from the feature covariance matrix, Spearman correlation generally requires additional information about the joint distribution. We therefore estimated Spearman correlation from resampled feature pairs combining the theoretical response predictions with calibration-derived realization errors. We sampled one predicted conditional mean and one complete calibration residual independently, uniformly and with replacement. We added the selected vectors and repeated this procedure 50,000 times to predict the Spearman correlation between *P* and *X*.**Supplementary Note S2.4** detailed this approximation and detection rates.

For the displayed high-gamma correction in **Fig. S2f**, we subtracted the predicted association from the held-out association:

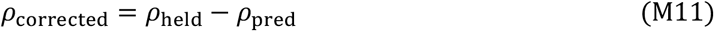

Here ρ_held_ is the empirical held-out Spearman correlation.

Each of 20 groups per frequency band contained 64 calibration parameter settings, with eight independent realizations per setting. FOOOF and IRASA-SPLIT received the same generated signals and were calibrated separately. Held-out evaluation used 128 input-parameter settings sampled independently of calibration in each group, with four independent realizations per setting. Calibration and held-out trials were sampled independently using the rule as described under *Simulated signal generation*.

### Computational models

#### Two-dimensional dynamical models

Model structure and inputs: We asked whether a minimal two-dimensional ordinary differential equation system can induce output feature correlation and whether weak nonlinear dynamics preserve that prediction. The linear model was

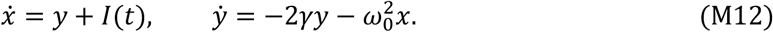

Here *x* is the measured state, *y* is an auxiliary state, *I*(*t*) is input, *γ* > 0 is the damping rate and *ω*_0_ is angular frequency. Eigenvalues are real for *γ*/*ω*_0_ > 1 and complex for *γ*/*ω*_0_ < 1. The nonlinear model was the cubic Hopf normal form,

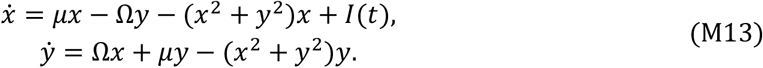

The rate *µ* < 0 stabilizes the origin; Ω is angular rotation frequency.

Population validations fixed *ω*_0_/(2π) = Ω/(2π) = 25 Hz. The stable-node regime fixed *γ*/*ω*_0_ = 2 for the linear model (**Fig. S3b**) and *µ*/(2π) = −40 Hz for nonlinear model (**Fig. S4b**). The latter is strongly damped but still has complex eigenvalues *µ* ± *i*Ω. **Supplementary Note S4** gave the eigenvalues and transfer functions.

For stable-node regime in **Fig. S3b** and strongly damped condition in **Fig. S4b**, trials varied the shape of a bump-plus-aperiodic input with target PSD ,*Q*(*f*; **θ**):

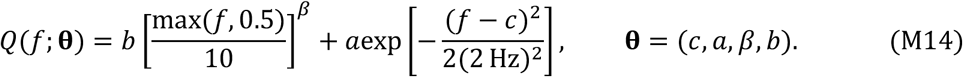

Here *c* is bump centre, *a* its height, *β* background colour and *b* background amplitude. We used 1,000 equally spaced target correlations *ρ*_0_ ∈ [−0.995,0.995]. For each condition, two centred, unit-variance, orthogonal Gaussian vectors *u, v* of length 64 defined 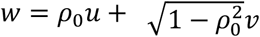. Trial parameters were (*c, a, β, b*) = (10 + 2.2*u*, 6 + 0.7*u*, −0.5 + 0.32*w*, 1 + 0.12*w*), clipped respectively to [5.5,18], [1,12], [-1.4,0.5] and [0.3,2]. In simulation, random-phase inverse Fourier synthesis generated real inputs in time domain with the shape in equation (M16). The zero-frequency component was zero, negative frequencies were conjugate symmetric, and each coloured trace was demeaned and normalized to RMS 0.02.

For resonant conditions with stable-complex eigenvalues in for **Fig. S3e**,**f** and **Fig. S4e**,**f**, input had no imposed oscillatory peak:

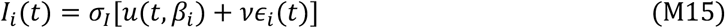

Where *u*(*t, β*_*i*_) is colored noise of RMS σ_*I*_ = 0.02 with slope 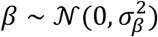 truncated to [-0.8,0.8], and realized with random phase. ϵ_*i*_(*t*) is zero-mean, unit-variance Gaussian white noise. σ_β_ and white-noise ratio ν were fixed within a condition. A three-dimensional Latin-hypercube design sampled (*γ*/*ω*_0_, σ_β_, ν) in linear model or (*µ*/2π, σ_β_, ν) in nonlinear model. The first 1,000 accepted conditions each contained 256 trials. Ranges, selection and the frequency normalization of the coloured component are specified in **Supplementary Note S4**.

Theoretical prediction: Let *J* be the Jacobian at the unforced fixed point, **b** the input vector and **v** the output-selector vector, which selects *x* in the two-dimensional systems. The scalar transfer and predicted output PSD are

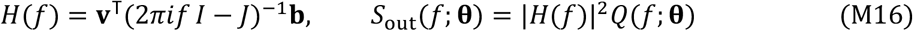

Here the symbol *I* is the identity matrix. For the lienar model this transfer is exact; for the nonlinear model it is the first-order approximation in state amplitude. If *G* contains the derivatives of (*P, A*) with respect to input generating parameters **θ** = (*c, a, β, b*), then

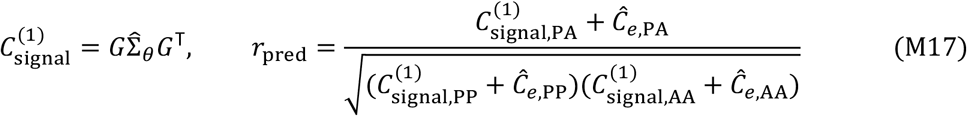

Here 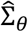 is the sample covariance matrix of the actual trial parameters and Ĉ_*e*_ is calibration-derived realization covariance. To obtain *G*, we used centred finite differences of the feature map (*c, a, β, b*) ↦ (*P, A*) derived from the transfer function. Bump-input derivatives were evaluated at (*c, a, β, b*) = (10,6, −0.5,1); aperiodic-only derivatives were evaluated at *β* = 0. Peak and slope derivative formulas are given in **Supplementary Note S5**. We obtain Ĉ_*e*_ from calibration trials (see below).

Simulation, calibration and statistical analysis: In simulation, we prepended the final 2.048 s of the trace as burn-in. Simulations of the nonlinear model integrated the complete cubic equations with explicit Heun steps of 0.25 ms; analysis windows were 32.768 s for bump-driven and 131.072 s for aperiodic-only conditions. Linear model simulation used a finite-window observation model.

For directly simulated outputs in nonlinear model, Welch spectra used 4.096-s Hann segments, 50% overlap and constant detrending. We smoothed log power with a Gaussian of standard deviation 0.9765625 Hz. A cubic spline of log power *s*(*f*) identified the largest interior maximum in the peak band *P*:

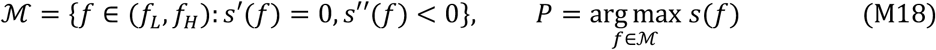

The signed aperiodic slope *A* was estimated by ordinary least squares (OLS) regression of log power on log frequency over ℱ_*A*_:

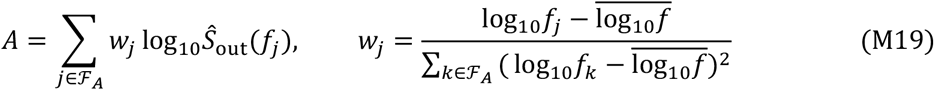

The set ℱ_*A*_ contained bins from 100 to 1,000 Hz; peak searches covered 5-100 Hz. Boundary or absent peaks were invalid. **Supplementary Note S6** records the additional validity masks used by each implementation.

The calibration model perturbed the theoretical PSD with unit-mean Gamma noise, extracted features by three-point peak interpolation and slope fitting, and generated repeated Gaussian feature errors with the resulting covariance. Within-trial centring across three calibration realizations estimated Ĉ_*e*_ by equation (M9)., but corrected with degree of freedom (See **Supplementary Note S2.3**). **Supplementary Note S6** specifies the Gamma parameters and sample counts.

For each frozen trial ensemble, evaluation averaged Pearson correlations calculated separately in five independent held-out realizations. Outcomes and theory used their recorded validity masks. Trial clouds in **Fig. S3f2** and **Fig. S4f2** pooled across realizations were descriptive; their phase alignment and effective units are specified in **Supplementary Note S6**.

#### Whole-brain excitatory-inhibitory model

Model structure: We used the 55-area marmoset cortical model, anatomical connectivity, hierarchical gradients and parameterization of Li, Li and Wang.^4^ Each area contained excitatory (E) and inhibitory (I) rate populations. In compact form,

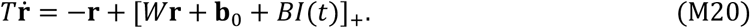

The vector **r** contains firing rates of all 110 rate populations (E group and I group for each of 55 areas), *T* is the diagonal matrix of population time constants, *W* contains gain-scaled local and long-range weights, and **b**_0_ is constant background drive. The column *B* has a single entry one at the stimulated E population and zeros elsewhere. Rectification [*z*] _+_ = *ma*x(*z*, 0) acts componentwise. Blocks and parameters are in **Supplementary Note S7**. The baseline background satisfies **b**_0_ = **r**_0_ – *W***r**_0_, with E-I rates of 10 and 35 Hz.

Stable-node input-output map: We asked how input feature association is transferred through a fixed resting network. Connectivity, background, input area V1 and output area PG were fixed. The input had equation (M16)’s form with four generating parameters **θ** = (*c, a, β, b*).

For the theoretical prediction of *P* − *A* correlation, we solved the fixed point **r**^*^ = [*W***r**^*^ + **b**_0_] _+_ and formed

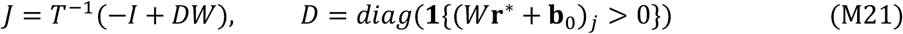

Here, 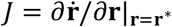 is the Jacobian of the network dynamics evaluated at the fixed point. Transfer predictions used equation (M16) with effective input vector *T*^−1^*DB*:

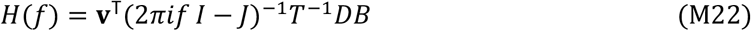

Background and active-set conventions are detailed in **Supplementary Note S7**. Then we used equation (M19) to calculate the prediction of *P* − *A* correlation *r*_pred_, evaluated around the baseline parameters(*c, a, β, b*) = (10,6, −0.5,1). Calibration pooled three repeated realizations of each trial from 14 independent 64-trial input ensembles. The main 1,000-condition comparison in **Fig. 2b** used one evaluation realization per condition, and the separate zero-input-association control in **Fig. 2c** used five. The design families and derivative steps are specified in **Supplementary Note S8**.

In **Fig.2b**, each of 1,000 conditions used a different target association and 64 trials. (*c, a, β, b*) is constructed as *c*=10+0.6*u, a*=6, *β*=-0.5+0.16*w* and *b*=1, where *u,w* were constructed as in “Model structure and inputs” under “Two-dimensional dynamical models”.

For the separate zero-input-association control in **Fig. 2c**, 100 independently constructed 64-trial ensembles had extracted input *P* − *A* correlation numerically zero, obtained by iterative orthogonalization of the input coordinates. The exact input normalization and validity-conditioned orthogonalization are specified in **Supplementary Note S8**.

Simulation used the full 110-state Heun integration with stepsize of 0.25 ms, implemented by finite causal convolution on the fixed active branch. Direct rectified pilot simulations checked numerical agreement on selected trajectories (**Supplementary Note S6**). The analysis window was 32.768 s after 2.048-s burn-in. Output extraction followed (M20)-(M21), with a 5-20-Hz peak band.

Stable-complex network prediction: We asked whether aperiodic-only drive produces correlated output features when the network is in a stable complex mode achieved by tuning PG inhibitory-to-excitatory coupling multiplier S. The input had equation (M17)’s form with three generating parameters θ=(*σ*_*I*_,*β,ν*).

For each network condition, we calculated fixed point, active set, Jacobian and transfer function as in (M21)-(M22). We propagated each trial’s input spectrum through the full-network transfer and accounted for the finite Hann-window kernel,. Extracting peak frequency *P* and aperiodic exponent *X* = −*A* according to (M18)-(M19) and expanding it to first order in the trial level input-colour parameter *β* at *β*_0_ = 0 as in (M3)-(M4) gave a deterministic feature pair **m**_*i*_ = (*P*_*i*_, *A*_*i*_)^⊤^. The kernel and parameter fitting process are specified in **Supplementary Note S8.2**. The predicted correlation was

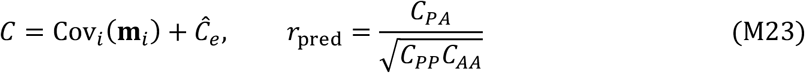

Here Cov_*i*_(**m**_*i*_) is the sample covariance across trials, and Ĉ_*e*_ is the condition-specific realization-error covariance calculated from calibration realizations. Each realization regenerated random phases and white-noise samples while keeping trial parameters fixed. Calibration centred repeated output feature pairs within each trial and pooled their residual outer products, normalized by the total within-trial degrees of freedom. Calibration and held-out realizations used independent random time series.

We used the PG excitatory population for both input and readout. Within each condition, we fixed the PG inhibitory-to-excitatory coupling multiplier *S* (range: 3.2662-6.6938), input colour spread σ _β_ (range: 0.05022-0.41095) and white-noise ratio ν (range: 0.07207-0.49997). Across trials, *β* followed a zero-mean Gaussian distribution with standard deviation σ _β_, truncated to [−0.8,0.8]. The coloured input had RMS 0.02 Hz, and the independent white component had standard deviation 0.02ν Hz. The retained ensemble contained 1,000 conditions with 256 trials each. Each condition reused its trial parameters across three calibration and five held-out realizations. Latin-hypercube design sampled (*S*, σ _β_, ν) for each condition with additional enriched candidates; sampling and acceptance are specified in **Supplementary Note S8.2**.

Calibration and held-out responses used the full-network fixed-branch Heun convolution with a 0.25-ms step (**Supplementary Note S6**). Analysis used 131.072-s records after a 2.048-s burn-in. We estimated the output PSD using Welch’s method. Subsequent feature extraction followed (M18)-(M19) with a 5-100-Hz peak band. We calculated the Pearson correlation across trials separately for each held-out realization and averaged the five correlations to give the simulated correlation.

Structural disconnection: We asked how long-range disconnection changes output correlation under matched stimulation. For a disconnected-area set *D*_*S*_, the theoretical effect was

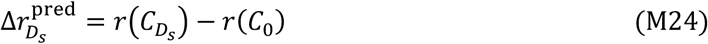

Here r(C_0_) and 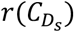 denote the *P* − *A* correlation in intact and node-deleted network, respectively. We computed each structure’s fixed point, active set, Jacobian and transfer function according to (M23)-(M24). We extracted peak-frequency and signed-slope pairs 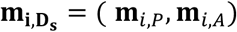 from the output PSD derived by transfer function separately for each trial. Their covariance was combined with structure-specific calibration:

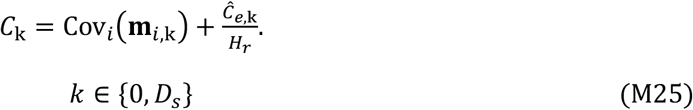

Calibration estimated single-realization covariance 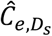 according to (M9) with a correction of degree of freedom (See **Supplementary Note S2.3**). Here *H*_*r*_ counts the number of held-out spectra averaged within each trial before feature extraction.

We zeroed all incoming and outgoing long-range weights incident on *D*_*S*_, retaining local E-I circuits and all network state variables. Background drive, trial parameters and paired random seeds were held fixed across intact and disconnected networks. Input was constructed according to (M16) and delivered to A6Va. Its four parameters (*c, a, β, b*) were drawn around (10,6, −0.5,1) using the multivariate Gaussian construction in **Supplementary Note S8.3**. The standardized perturbation covariance was 0.3^2^Σ_*d*_; Σ_*d*_ had condition number 100 before sampling and clipping. Coordinate scales, bounds and the covariance matrix are specified in **Supplementary Note S8**.

The theory-simulation scatter in **Fig.2i** and the node-count comparison in **Fig. 2j** used 82 deletion patterns and 128 trials per pattern, with three calibration and five held-out realizations. The node-count comparison in **Fig.2j** used 60 patterns, 12 for each node deletion count from one to five, with 512 trials per pattern, two calibration and two held-out realizations for each trial. Deleted outputs and outputs inactive at the modified fixed point were excluded. An eligible output was direct if it received a positive intact-network edge from at least one disconnected area; otherwise, it was non-direct. Pattern selection is detailed in **Supplementary Note S8.3**.

Each trajectory started at its own network structure’s fixed point. Held-out trajectories integrated all 110 rectified equations directly with Heun. The integration step was 0.25 ms and burn-in was 2.048 s. Analysis windows were 32.768 s for **Fig.2i2** and 16.384 s for **Fig.2i1**. Feature extraction followed (M20)-(M21). Within each trial, we averaged held-out PSDs before extracting one feature pair, then calculated the correlation across trials for each structure. The simulated effect was 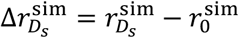.

#### Empirical tests of model predictions

##### Transcranial electrical stimulation

tES was delivered through scalp pad electrodes positioned over frontal and occipital sites while sEEG activity was recorded simultaneously (**Fig. 3a**). We included all valid sEEG contacts (following pre-processing procedure) since tES affected all contacts.

Stimulation comprised three aperiodic waveforms (white, pink and brown noise) and four periodic sinusoidal waveforms (8, 32, 64 and 128 Hz). Aperiodic stimulation was used to test changes in oscillatory activity, and periodic stimulation to test changes in the aperiodic component. Each stimulating condition had one trial and was randomly presented. In each trial, stimulation lasted 30 s and its amplitude increased linearly from 0 to 0.5 mA over the first 10 s and remained at 0.5 mA for the next 20 s. Consecutive trials were separated by a 30-s stimulation-free interval to minimize carryover effects. Because concurrent stimulation may produce residual spectral flattening at higher frequencies, we did not apply direct artifact subtraction, which could distort the underlying spectral profile. Instead, analyses were restricted to frequencies below 30 Hz, outside the affected high-frequency range.

Effects of aperiodic stimulation on oscillatory peaks: Periodic peaks were extracted using the fitting procedure above and assigned to delta-theta (1-8 Hz), alpha (8-13 Hz) and beta (13-30 Hz). For each channel, time window and band, the highest-power peak was retained. Peak center frequencies were compared between the pre-stimulation baseline (-20 to 0 s) and steady-state stimulation (10-30 s), separately for white-, pink- and brown-noise stimulation. Times were relative to stimulation onset.

Effects of periodic stimulation on the aperiodic exponent: For stimulation at 8, 32, 64 and 128 Hz, IRASA fractal spectra were parameterized over 0.2-30 Hz using SPLIT. Analyses retained spectra best fitted by a single-knee model. The aperiodic exponent was compared between baseline (-20 to 0 s) and steady-state stimulation (10-30 s).

#### Inter-, pre-ictal, and ictal sEEG analysis

Expert epileptologists (X.H., D.L., and J.C.) annotated interictal, preictal and ictal periods and identified seizure-onset-zone (SOZ) channels by visual inspection and clinical electrophysiological criteria. The interictal baseline was a continuous, artifact-free 20-min segment distant from seizures. The preictal period comprised the 5 min immediately preceding electrographic seizure onset. To measure excitatory-inhibitory ratio for each contact and state, we used both time-resolved spectral exponent^16^ and fEI (functional excitation-inhibition ratio)^32^ as approximates. Aperiodic-periodic coupling was quantified as above and compared between states and between SOZ and non-SOZ channels.

#### CCEP mapping and RF-TC analysis

CCEPs were recorded during routine clinical single-pulse stimulation mapping. Square-wave pulses were delivered sequentially to adjacent contact pairs along each electrode shaft, advancing one contact at a time. Responses to stimulation at sites subsequently treated with RF-TC were identified retrospectively. CCEPs at other recording contacts were epoched from -0.5 to 0.8 s relative to stimulation onset and averaged across pulses, with a baseline of -200 to -20 ms. For each stimulation site, a recording contact was considered responsive if the averaged waveform exceeded ±6 baseline standard deviations in both an early window (11-30 ms) and a late window (81-250 ms), with the same polarity in both windows.

Contacts directly treated with RF-TC and bipolar channels containing these contacts were excluded. Recording contacts on the same electrode shaft as a stimulated RF-TC site were also excluded if they exhibited a qualifying CCEP response to that site. Each retained recording contact was assigned to a single group: CCEP+ if it exhibited a qualifying response to at least one RF-TC site, or CCEP-if it exhibited no qualifying response to any RF-TC site. Aperiodic-periodic coupling was quantified in 30-min sEEG recordings acquired 2-4 days before and 2-4 days after RF-TC, with timing varying across patients. Before-after changes in coupling were assessed separately within CCEP+ and CCEP-contacts, and the magnitude of change was compared between the two groups, as described below.

### Statistical analysis

Statistical analyses of the experimental data used two-sided Spearman correlation tests and linear mixed-effects models. Multiple comparisons were corrected using the Benjamini– Hochberg false discovery rate procedure, with statistical significance defined as an FDR-adjusted p < 0.05.

#### Linear mixed-effects models

Models were fitted using MATLAB’s fitlme function. In the wakefulness dataset, coupling coefficients were tested against zero separately in each of the five frequency bands using the intercept-only model:

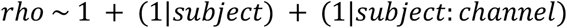

For aperiodic tES, peak center frequency was modelled as:

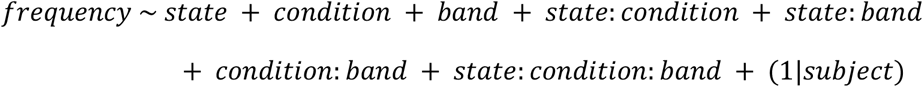

State contrasted baseline (-20 to 0 s) with stimulation (10-30 s); condition indexed white-, pink- and brown-noise stimulation; and band indexed delta-theta, alpha and beta.

For periodic tES, the post-knee exponent was modelled as:

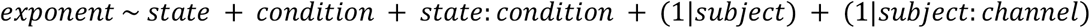

Condition indexed stimulation at 8, 32, 64 and 128 Hz, with the same state definition.

For interictal, preictal and ictal recordings, coupling was analyzed separately in each frequency band using the following model:

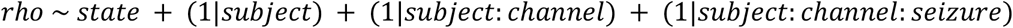

State denoted the recording period; preictal and ictal recordings were each compared with the interictal baseline. Changes in fEI and aperiodic exponent across recording periods were also assessed using linear mixed-effects models.

For the CCEP mapping and RF-TC analyses, pre- and post-RF-TC coupling coefficients were compared separately within CCEP+ and CCEP-contacts in each of the five frequency bands using the model:

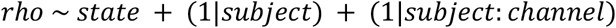

State distinguished recordings obtained before and after RF-TC. Random intercepts were included for participants and channels nested within participants to account for repeated measurements. To compare the magnitude of change between CCEP+ and CCEP− contacts, we calculated the absolute difference between post- and pre-RF-TC coupling coefficients for each contact, |Δrho| = |rho_post_ – rho_pre_|. Absolute changes were analyzed separately in each frequency band using the following model, with CCEP group (CCEP+ versus CCEP-) as the fixed-effect predictor.

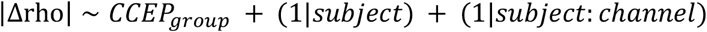

## Data and code availability

De-identified sEEG data are available upon request to the lead author. Access to raw clinical recordings requires a data-sharing agreement and institutional approval. Analysis code and the MATLAB and Python implementations of SPLIT are available at https://github.com/yu-han-lu/aperiodic-periodic-coupling.

## Acknowledgements

This study is supported by the National Natural Science Foundation of China 32271101, the Program of AI-Driven Initiative to Promote Research Paradigm Reform and Empower Disciplinary Advancement by Shanghai Municipal Education Commission (SMEC), Program of Introducing Talents of Discipline to Universities, Base B16018, and NYU Shanghai Boost Fund (X.T.). The Brain Science and Brain-like Intelligence Technology - National Science and Technology Major Project No. 2022ZD0204600 and the Natural Science Foundation of China Grant No. 31771147 (Z.L. and L.T.), Z.X.: Talent Program by SMEC, and boost fund by NYUSH.

## Author contributions

Y.L. and X.T. contributed to conceptualization; All authors designed the study; Z.L., L.T., Z.X., and Y.L. contributed mathematical modelling; D.L., X.H., and J.C. collected data; Y.L. and H.M. conceived of and coded the algorithm and analysed data; L.T., Z.X. and X.T. provided overall supervision and guidance throughout the project; all authors contributed to the manuscript.

