## Supplementary figures and tables for "State-dependent emergence of coupling between oscillatory and aperiodic neural activity"

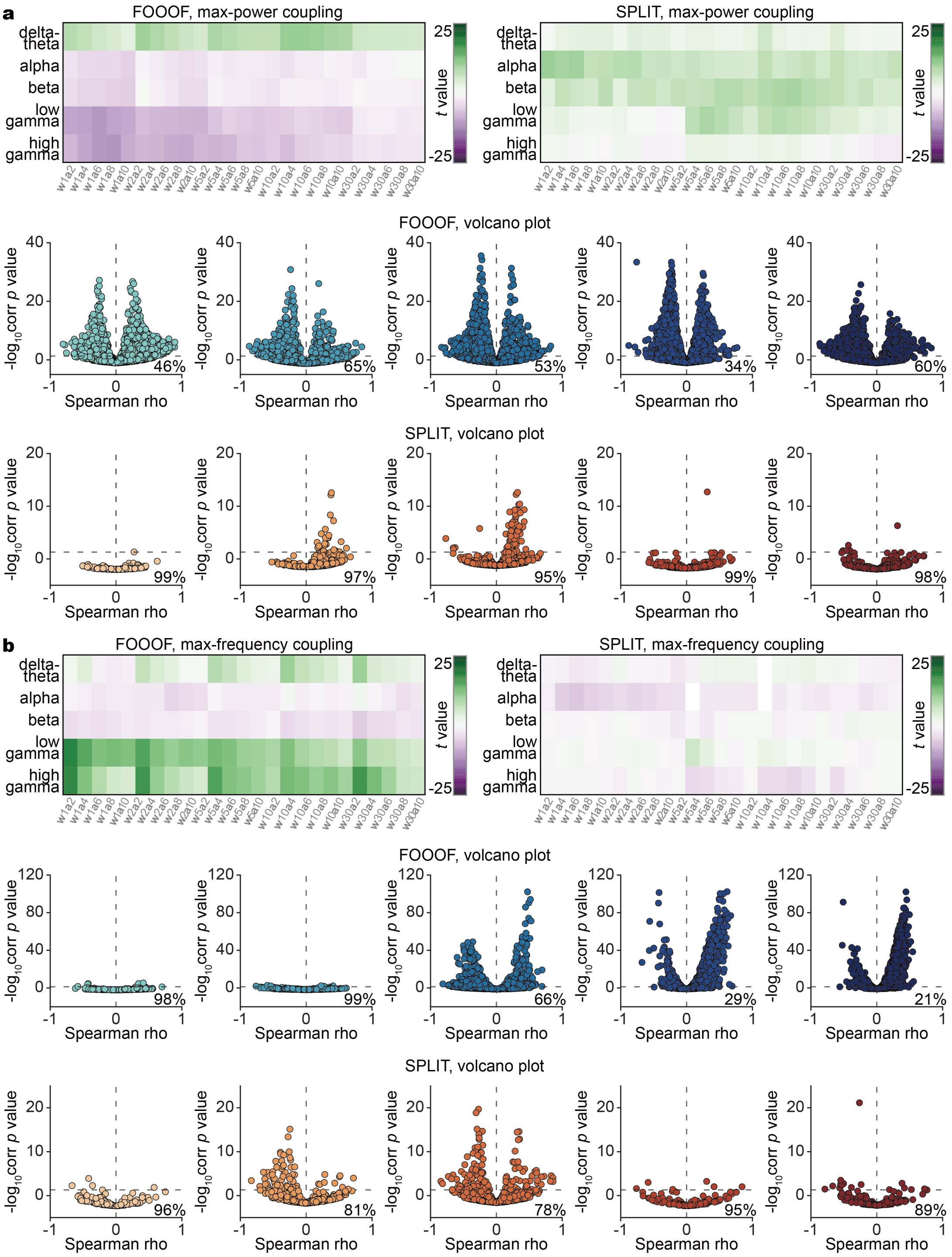


**Supplementary Figure 1 | Correlations of aperiodic exponent with alternative oscillatory peak measures.**

Awake sEEG recordings from 61 subjects were analysed using FOOOF and SPLIT. **a,** Within-contact Spearman correlations between aperiodic exponent and the power of the highest-power peak in each frequency band. **b,** Corresponding correlations between aperiodic exponent and the highest detected peak frequency. In both **a** and **b**, heatmaps show population-level *t* statistics across 25 combinations of spectral-window duration (*w* = 1, 2, 5, 10 or 30 s) and temporal averaging width (*a* = 2, 4, 6, 8 or 10 neighbouring spectra). Scatter plots below the heatmaps show contact-level correlations and their significance for 6-s window length (50% overlap), averaging 6 neighbouring spectra, with FOOOF above and SPLIT below. Columns show delta-theta, alpha, beta, low gamma and high gamma, from left to right. Each point represents one contact. Dashed lines mark zero correlation and the significance threshold. Percentages indicate contacts without significant correlations.

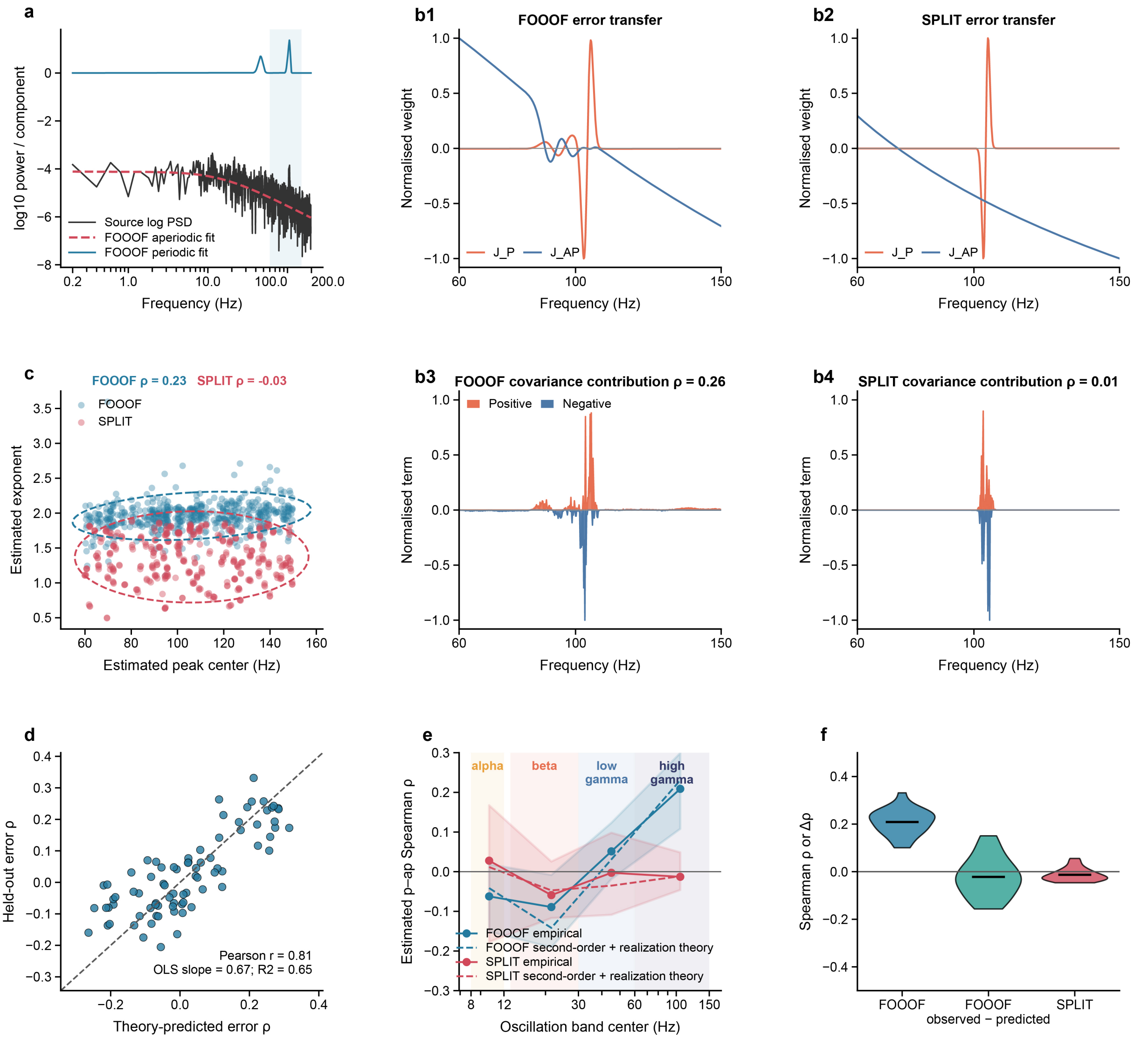


**Supplementary Figure 2 | Error propagation predicts apparent periodic-aperiodic coupling in simulated signals.**

Synthetic signals combined an Ornstein–Uhlenbeck background with an independently generated narrowband component. Predictions were derived from calibration signals and tested on independently generated signals. Calibration and held-out parameter settings were sampled independently from the same band-specific distributions. Within each dataset, FOOOF and IRASA-SPLIT were applied to the same generated signals. Settings were weighted equally within each estimator, but estimator-specific validity masks were retained. **a,** Representative high-gamma power spectral density (PSD), with FOOOF aperiodic and periodic fits. **b1,b2,** Normalized local weights showing how a change in log power at each frequency affects estimated peak frequency and aperiodic exponent for FOOOF and SPLIT. **b3,b4,** Corresponding first-order contributions to covariance between the two estimates. The reported theoretical correlations also include second-order responses to the generating parameters and variability across repeated signals with the same parameters. **c,** Estimated exponent versus peak frequency in a representative test group. Each point represents one simulated signal. Dashed ellipses show predicted covariance contours centred on the observed means. **d,** Predicted versus observed correlations between FOOOF estimates across 80 groups, comprising 20 groups in each of four frequency bands. Each point represents one group, and the dashed line indicates equality. **e,** Observed and predicted correlations across alpha, beta, low gamma and high gamma. Solid lines and points show observed group means, dashed lines show predicted means, and shading spans the 2.5th to 97.5th percentiles of observed group correlations. **f,** High-gamma correlations for FOOOF and SPLIT, alongside the difference between observed and predicted FOOOF correlations ($\Delta\rho =$

$\rho_{observed} - \rho_{predicted}$.) Violins show distributions across 20 groups, with horizontal bars indicating means. Subtracting the predicted association reduced the mean FOOOF correlation towards zero. Observed and predicted feature correlations use Spearman's ρ. Prediction accuracy in **d** is summarized by Pearson correlation. Reported *P* values in **c** and **d** are from two-sided Spearman and Pearson correlation tests, respectively.

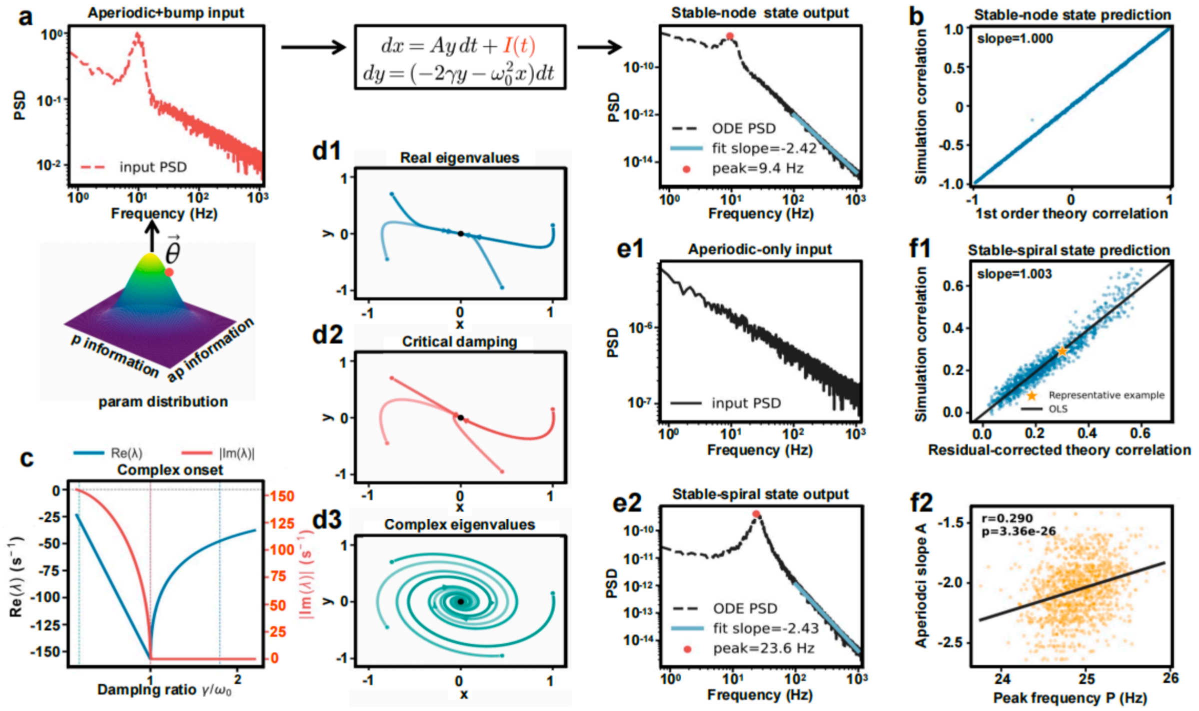


**Supplementary Figure 3 | First-order theory predicts coupling in a linear damped oscillator.**

Coupling is the Pearson correlation between output peak frequency *P* and signed aperiodic slope *A* across trials. Here *A* is the log-log spectral slope. **a,** Model framework and example input and output PSDs for an input containing an oscillatory peak and an aperiodic background. **b,** First-order predictions versus finite-window simulations using the exact linear frequency response in the overdamped regime ($\gamma/\omega_0 = 2$). Each point represents one of 1,000 input conditions, each containing 64 trials. The diagonal line indicates equality. **c,** Real and absolute imaginary parts of the eigenvalues as the damping ratio $\gamma/\omega_0$ varies. Critical damping occurs at $\gamma/\omega_0 = 1$. **d1–d3,** Trajectories from direct integration in the overdamped, critically damped and underdamped regimes, respectively. **e1,e2,** Aperiodic-only input and the corresponding underdamped output ($\gamma/\omega_0 = 0.16$), showing an output peak without an imposed input peak. Cyan lines in **a** and **e2** show slope fits, and red dots mark spectral peaks. **f1,** Predictions incorporating calibration-derived covariance from repeated realizations versus independent finite-window simulations using the exact linear frequency response. Each point represents one of 1,000 accepted conditions, each containing 256 trials. Simulated correlations were averaged across five realizations. The line shows an ordinary least-squares fit, and the star identifies the condition in **f2**. **f2,** Output peak frequency and signed slope for 256 trial settings across five realizations, pooled after alignment within each realization. Points and the fitted line describe their association.

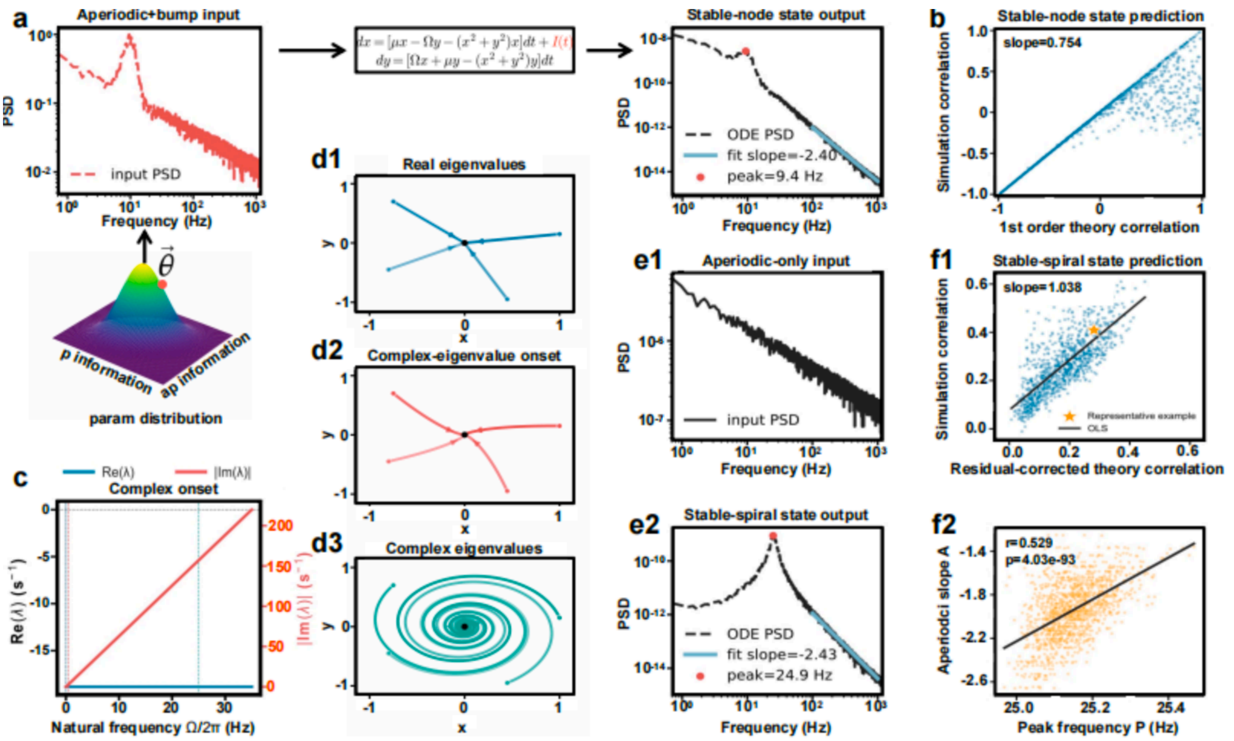

**Supplementary Figure 4 | First-order theory approximates coupling in a nonlinear dynamical model.**
Coupling is the Pearson correlation between output peak frequency $P$ and signed aperiodic slope $A$ across trials, with aperiodic exponent $\chi = -A$. **a,** Model framework and example input and output PSDs for the cubic Hopf normal form driven by an oscillatory peak and an aperiodic background. **b,** First-order predictions versus simulations of the full nonlinear equations in a strongly damped regime. Each point represents one of 1,000 input conditions, each containing 64 trials. The diagonal line indicates equality. **c,** Real and absolute imaginary parts of the eigenvalues as $\Omega/(2\pi)$ varies at fixed $\mu < 0$. Their real parts remain negative, so the equilibrium remains stable. **d1–d3,** Representative trajectories at $\Omega = 0$ and two positive values of $\Omega$, showing decay towards the equilibrium with increasing rotation. This marks the introduction of rotational dynamics, not a loss of equilibrium stability or a Hopf bifurcation. **e1,e2,** Aperiodic-only input and the corresponding output in a weakly damped regime, showing an output peak without an imposed input peak. Cyan lines in **a** and **e2** show slope fits, and red dots mark spectral peaks. **f1,** Predictions incorporating calibration-derived covariance from repeated realizations versus independent nonlinear simulations. Each point represents one of 1,000 accepted conditions, each containing 256 trials. Simulated correlations were averaged across five realizations. The line shows an ordinary least-squares fit, and the star identifies the condition in **f2**. **f2,** Output peak frequency and signed slope for 256 trial settings across five realizations, pooled after alignment within each realization. Points and the fitted line describe their association.

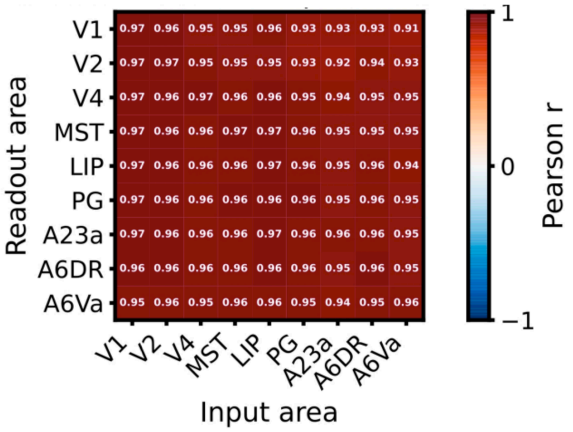


**Supplementary Figure 5 | Correspondence between predicted and simulated coupling across cortical pathways.**
The baseline analysis in Fig. 2b was extended to all 81 combinations of nine input and nine output areas in the 55-area excitatory-inhibitory network. Input was delivered to the excitatory population of the area indicated by each column, and output was measured from the excitatory population of the area indicated by each row. Each cell shows the mean held-out Pearson correlation $r(P, \chi)$ across five nonlinear simulation realizations for one registered 64-trial rank-one input ensemble. Trials were retained only when peak-frequency and exponent estimates were finite and the peak was an unambiguous interior maximum in all three calibration and five held-out realizations. The common-valid count ranged from 21 to 64 trials (median, 60). Colours and numerical labels indicate the same correlation coefficient.

**Table S1.** Leave-k-out sensitivity of the beta-band state effect

| Patients excluded (k) | Patients remaining | Combinations tested | Combinations remaining significant | Proportion (%) |
|---|---|---|---|---|
| 1 | 18 | 19 | 19/19 | 100.00% |
| 2 | 17 | 171 | 171/171 | 100.00% |
| 3 | 16 | 969 | 964/969 | 99.50% |
| 4 | 15 | 3876 | 3791/3876 | 97.80% |
| 5 | 14 | 11628 | 11050/11628 | 95.00% |

Linear mixed-effects models were refit after excluding every unique combination of k patients ($N$ = 19). The ictal versus interictal coefficient remained significant ($p < 0.05$; FDR corrected) in the tabulated fraction of combinations.

**Table S2.** Demographic and clinical characteristics by dataset

| Dataset | *n* | Age (y) | Sex (F/M) | Contacts analysed |
|---|---|---|---|---|
| Wakefulness | 61 | 21.4 ± 11.8 | 25 / 36 | 52 (17–95) |
| Simultaneous tES | 27 | 22.1 ± 8.7 | 11 / 16 | 68 (25–102) |
| Interictal, preictal and ictal | 19 | 23.2 ± 10.8 | 7 / 12 | 59 (34–90) |
| RF-TC | 15 | 27.5 ± 8.2 | 10 / 5 | 128 (101–206) |

Age is reported as mean ± SD (years). Sex is reported as number of female / male patients. Contacts analysed are reported as median (range) per patient. tES, transcranial electrical stimulation; RF-TC, radiofrequency thermocoagulation.
