## Supplementary notes for "State-dependent emergence of coupling between oscillatory and aperiodic neural activity"

### Supplementary Notes
#### S0 Analysis map and notation

**Supplementary Note Table 1 | Theory prediction realization and simulation methods used in figures.**

| **Analysis and panels** | **Simulation or model level** | **Theory map** | **Realization term used by theory** | **Sections in Supplementary Notes** |
|---|---|---|---|---|
| Estimator validation, **Fig. S2a–f** | 30-s time-domain OU-plus-oscillation records fitted separately by FOOOF and simulation-specific IRASA–SPLIT | Quadratic response map in $(c, w, q, f_k)$ for panels c–f; local log-spectral influence rows only for panel b | Complete calibration residual pairs are resampled to predict Spearman correlation; independent held-out time series provide the comparison | S1-3 |
| Nine-input-by-nine-readout pathways, **Fig. S5** | Full 55-area, 110-state network; population outcomes use direct rectified Euler integration | Path-specific full-network transfer followed by first-order input-parameter propagation | Path-specific prediction calibration and validity masks; retained denominators range from 21 to 64 trials | S8.1 |
| Linear two-dimensional model, **Fig. S3** | Exact two-state linear dynamics | First-order generating-parameter map through the exact resolvent | Finite-window observation covariance generated by the specified Gamma spectral and feature-error model | S4-6 |
| Cubic Hopf model, **Fig. S4** | Direct two-state nonlinear cubic Heun integration | First-order parameter map obtained from the transfer at the origin | Finite-window observation covariance generated by the specified Gamma spectral and feature-error model | S4-6 |
| Resting stable-node network, **Main Fig. 2b,c** | Full 110-state fixed-active-branch discrete Heun convolution | First-order feature map $G\Sigma_{\theta}G^{\top}$ about the resting reference | Degrees-of-freedom-corrected single-realization covariance from three realizations of 14 independent 64-trial calibration ensembles | S7, S8.1 |
| Stable-complex network, **Main Fig. 2d–g** | Full 110-state fixed-active-branch Heun calculation with a condition-specific fixed point and transfer | First-order expansion of the transfer-derived feature mao about aperiodic slope $\beta = 0$ | Condition-specific single-realization covariance from three calibration realizations; five held-out realization-wise correlations form the comparator | S7, S8.2 |
| Structural disconnection, **Main Fig. 2h-j** | Direct 110-state rectified Heun integration for held-out outcomes, with intact and deleted structures solved separately | Difference between two structure-specific complete trial-wise feature maps | Structure-specific single-realization covariance scaled by the number of PSDs averaged within each trial | S7,S8.3 |

**Supplementary Note Table 2 | Notation and sampling units.**

| Symbol or unit | Meaning |
|---|---|
| $P$, $\chi$, $A$ | Peak centre in Hz; positive fitted aperiodic exponent; signed log–log slope. For the same slope definition, $A = -\chi$. |
| $\boldsymbol{\theta}$ | OU: $(c, w, q, f_k)^{\mathsf{T}}$. Bump-driven dynamical models: $(c, a, \beta, b)^{\mathsf{T}}$. |
| $g_a$, $H_a$ | Gradient and Hessian of the conditional mean of feature $a$ with respect to generating parameters. |
| $J_p$, $J_{ap}$ | Local sensitivity of estimated peak frequency and exponent to log-spectral perturbations. |
| $J$, $H(f)$ | State-space Jacobian at a fixed point; input-to-output transfer function. |
| $C_m$, $C_e$ | Covariance of conditional feature means; covariance associated with repeated realizations at fixed parameters. |
| Condition or group | One experimental design defining a distribution of input parameters and, where varied, network or damping parameters. |
| Trial or parameter setting | One generating-parameter vector within a condition. |
| Realization | A new random time series or observation-model draw at the same trial parameters. |
| Calibration and held-out | Data used to estimate coefficients or realization variability; independent outcomes used to evaluate prediction. |

### S1 OU process simulation and spectral estimator inputs

Equations (M1)–(M2) in Methods define the four input-generating parameters, the OU recursion, the filtered oscillatory component, the composite signal, all parameter distributions and the record length. They are not restated here. This note derives the discrete OU spectrum, explains the finite-record normalization in M2 and gives implementation details needed to reproduce **Fig. S2**.

#### S1.1 Discrete OU process spectrum

Equation (M1) generates the aperiodic component with an Euler–Maruyama OU recursion at $\Delta t = 1$ ms and initializes the state at zero. The analyses in **Fig. S2a,c–f** use records generated by this recursion. Equations (S1)–(S3) below are analytic consequences of M1: they give the stationary variance, lag covariance and exact discrete spectrum for $a_0 = 1 - \Delta t/\tau$, with $\tau = (2\pi f_k)^{-1}$. They are not replacement simulation equations. Equation (S4) separately bounds the finite-record variance loss caused by the zero initial state and is not plotted in a panel.

Write the autoregressive coefficient in M1 as $a_0 = 1 - \Delta t/\tau$. For $|a_0| < 1$, a stationary variance exists. If $v = \mathrm{Var}(x_n)$, independence of $x_n$ and the next innovation gives the Yule–Walker equation and solution

$$v = a_0^2 v + \Delta t, \quad v = \frac{\Delta t}{1 - a_0^2}. \tag{S1}$$

Applying the same recursion across $|\ell|$ lags gives

$$R_x[\ell] = \mathrm{Cov}(x_n, x_{n+\ell}) = v \;\; a_0^{|\ell|}. \tag{S2}$$

The spectral density is the discrete-time Fourier transform of this covariance. Summing the two geometric series for positive and negative lags and multiplying by $\Delta t$ to express density per hertz gives

$$S_x^{\mathrm{disc}}(f) = \Delta t \sum_{\ell=-\infty}^{\infty} R_x\,[\ell] e^{-2\pi i f \ell \Delta t} = \frac{\Delta t^2}{1 + a_0^2 - 2a_0 \cos(2\pi f \Delta t)}. \tag{S3}$$

This is the stationary two-sided spectrum; the one-sided spectrum doubles positive bins except Nyquist. Expanding the denominator for small $\Delta t$ recovers a constant multiple of $[\tau^{-2} + (2\pi f)^2]^{-1}$. Thus the generator has the intended knee and an asymptotic exponent of two, but a fitted finite-band exponent can differ from two because it is estimated from a finite, knee-shaped discrete spectrum.

Equation (M1) initializes $x_0 = 0$, whereas (S3) assumes stationarity. For $x_n$, the missing variance relative to stationarity is $v a_0^{2(n+1)}$; averaging this loss across an $N$-sample record gives

$$\frac{1}{Nv} \sum_{n=0}^{N-1} v\, a_0^{2(n+1)} = \frac{a_0^2 (1 - a_0^{2N})}{N(1 - a_0^2)}. \tag{S4}$$

At the smallest sampled knee (approximately 0.499 Hz), this fraction is about 0.0053 and is smaller for faster knees. This bounds the initialization effect on record-averaged OU variance, but is not a sensitivity analysis of fitted correlations.

#### S1.2 Parameter sampling criteria

In **Fig. S2**, input PSD in each trial is determined by oscillation centre $c$,  passband width $w$, oscillatory-strength parameter $q$ and  knee frequency $f_k$. For alpha-, beta-, low-gamma- and high-gamma-band inputs, the oscillation centre $c$ was sampled uniformly from 8-12, 13-30, 30-60 and 60-150 Hz, respectively. In all bands, the passband width was sampled as $w \sim U(1.5,6)$ Hz. The oscillatory-strength parameter was sampled as $q \sim U(2.8,3.8)$ in the first three bands and $q \sim U(28,34)$ in the high-gamma band. The parameters $c$, $w$ and $q$ were sampled independently. The knee frequency $f_k$ was sampled by stratified inverse-CDF sampling from an empirical distribution containing 3,126 positive, finite knee values (i.e., empirical knee distribution in **Fig. 1**). We generated 32 random permutations of the sampled knee values and retained the assignment with the smallest absolute Spearman correlation $|\rho_S(c, f_k)|$, as stated in detail below.

##### S1.3 Stratified empirical knee assignment

The simulated groups in **Fig. S2c–f** draw $f_k$ from the empirical knee distribution. Methods states that the group design reduces accidental centre–knee association before simulation. Equation (S6) specifies that sampling and permutation step so that the calibration and held-out ensembles can be reconstructed without treating the empirical knees as independent uniform draws.

Methods gives the parameter ranges and ensemble sizes. The only additional sampling step was the finite-sample allocation of empirical knees. Let $F_k^{-1}$ be the linearly interpolated empirical quantile function:

$$p_i = \frac{i + U_i}{n}, \;\; U_i \sim U(0,1), \;\; i = 0, \ldots, n-1, \quad k_i = F_k^{-1}(p_i). \tag{S6}$$

Here, $n$ is the number of distinct generating-parameter settings, and hence knee-frequency draws, in the ensemble; $i = 0, \ldots, n-1$ indexes the equal-probability strata, $p_i \in [i/n, (i+1)/n]$ is the resulting quantile level, and $F_k^{-1}$ maps that level to the preliminary knee-frequency draw $k_i$.

Thrity-two random permutations of the $k_i$ were paired with the sampled centres, and the permutation minimizing the absolute Spearman correlation $|\rho_S(c, k)|$ was retained. This reduces incidental finite-sample centre–knee association. Calibration and held-out ensembles used independent draws and the same rule.

#### S2 Parameter-to-feature covariance and calibration

##### S2.1 Second-order covariance of the quadratic response map

The broad parameter distributions used for Fig. S2c–f require the quadratic response surface described in Equations (M3)–(M4) in Methods. The derivation below retains $H_a$ and $H_b$ and expresses the resulting feature covariance through the empirical second-, third- and fourth-order moments of the sampled parameter vectors.

The quadratic term has expectation $1/2\,\mathrm{tr}(H_a\Sigma)$. Subtracting this expectation from the second-order approximation gives

$$m_a^{(2)} - \mathbb{E}m_a^{(2)} = g_a^{\mathsf{T}}\mathbf{d} + 1/2\left(\mathbf{d}^{\mathsf{T}}H_a\mathbf{d} - \mathrm{tr}(H_a\Sigma)\right). \tag{S7}$$

Multiplying the centred expansions in (S7) for features $a$ and $b$ and summing over all indices gives $C_{ab}^{(2)} = Cov(m_a^{(2)}, m_b^{(2)})$ :

$$C_{ab}^{(2)} = \sum_{i,j} g_{a,i}\, \Sigma_{ij} g_{b,j}$$
$$+\frac{1}{2}\sum_{i,j,k}\left(g_{a,i}H_{b,jk} + g_{b,i}H_{a,jk}\right)T_{ijk}$$
$$+\frac{1}{4}\sum_{i,j,k,l} H_{a,ij}\, H_{b,kl}(U_{ijkl} - \Sigma_{ij}\Sigma_{kl}).$$

(S8)

#### S2.2 Count-weighted calibration and parameter fitting

The calibration in **Fig. S2c-f** fits feature responses from repeated simulations at each $(c, w, q, f_k)$ setting. Equations (S10)–(S11) make the implemented fit explicit.

To fit the conditional mean, we standardized each input parameter using its calibration mean and sample standard deviation,

$$x_{ij} = \frac{\theta_{ij} - \overline{\theta}_{C,j}}{s_{C,j}} \qquad \text{(S9)}$$

where $i$ indexes input settings and $j$ indexes the four generating parameters. These standardized values served as predictors in the regression of the within-setting mean feature estimates. For the second-order model, we fitted

$$\widehat{\boldsymbol{\beta}}_a = \underset{\boldsymbol{\beta}_a}{\operatorname{argmin}} \left[ \sum_{i:n_i \geq 2} n_i \left\{ \begin{array}{c} \overline{z}_{i,a} - \beta_{a,0} - \sum_{j=1}^{4} \beta_{a,j}\, x_{ij} \\ - \sum_{j=1}^{4} \beta_{a,jj}\, x_{ij}^2 - \sum_{j<k} \beta_{a,jk}\, x_{ij} x_{ik} \end{array} \right\}^2 + 10^{-4} \parallel \boldsymbol{\beta}_a \parallel^2 \right] \qquad \text{(S10)}$$

Here $\boldsymbol{\beta}_a$ contains the intercept and all linear, squared and pairwise interaction coefficients for feature $a$. After accounting for parameter standardization, these coefficients provide regression-based estimates of $g_a$ and $H_a$ in equation (M3). The weight $n_i$ is the number of valid realizations contributing to $\overline{z}_{i,a}$. The first-order fit retained only the intercept and linear terms.

At the calibration centre, conversion to original parameter units gives

$$\hat{g}_{a,j} = \frac{\beta_{a,j}}{s_{C,j}}, \quad \widehat{H}_{a,jj} = \frac{2\beta_{a,jj}}{s_{C,j}^2}, \quad \widehat{H}_{a,jk} = \frac{\beta_{a,jk}}{s_{C,j}s_{C,k}} \ \ (j \neq k). \qquad \text{(S11)}$$

Here, $s_{\mathrm{C,j}}$ denotes the sample standard deviation of the jth generating-parameter coordinate in the calibration set, used to express derivatives in original parameter units.

#### S2.3 Within-parameter realization covariance

Methods separates changes in the conditional feature mean from variability among repeated time series at fixed generating parameters. **Fig. S2c–f** combines both sources, as do the later two-dimensional and network predictions with analysis-specific normalizations. Equations (S12)–(S14) therefore derive the total covariance $C = C_m + C_e$, define the pooled and degrees-of-freedom denominators for $C_e$, and show how the predicted Pearson correlation is obtained from the resulting matrix.

Write $\mathbf{z} = \mathbf{m}(\boldsymbol{\theta}) + \mathbf{e}$, where the true conditional residual satisfies $\mathbb{E}(\mathbf{e} \mid \boldsymbol{\theta}) = 0$. To derive M6, centre this identity and expand its outer product. The mixed terms vanish because $\mathbb{E}[(\mathbf{m} - \mathbb{E}\mathbf{m})\mathbf{e}^{\mathsf{T}}] = \mathbb{E}_\theta[(\mathbf{m} - \mathbb{E}\mathbf{m})\mathbb{E}(\mathbf{e}^{\mathsf{T}} \mid \theta)] = 0$. Therefore

$$\mathrm{Cov}(\mathbf{z}) = \mathbb{E}[(\mathbf{m} - \mathbb{E}\mathbf{m})(\mathbf{m} - \mathbb{E}\mathbf{m})^{\mathsf{T}}] + \mathbb{E}(\mathbf{e}\mathbf{e}^{\mathsf{T}}) = C_m + C_e. \tag{S12}$$

For the fitted calibration pairs, $\mathbf{e}_{ih} = \mathbf{z}_{ih} - \bar{\mathbf{z}}_i$. Pooling these complete two-dimensional residuals preserves the association between peak and exponent errors. Let $N_e = \sum_{i:n_i \geq 2} n_i$ and $D_e = \sum_{i:n_i \geq 2} (n_i - 1)$. The two relevant normalizations are

$$\hat{C}_e^{\text{pool}} = \frac{1}{N_e} \sum_{i:n_i \geq 2} \sum_{h \in V_i} \mathbf{e}_{ih}\, \mathbf{e}_{ih}^{\mathsf{T}}, \quad \hat{C}_e^{\text{df}} = \frac{N_e}{D_e} \hat{C}_e^{\text{pool}}. \tag{S13}$$

The first is exactly the covariance of the empirical residual distribution sampled uniformly in (M8). The second corrects the loss of degrees of freedom caused by estimating a mean within each setting. For **Fig. S2c–f**, residual pairs were resampled without degrees-of-freedom rescaling, so the displayed predictions used $\hat{\mathrm{C}}_{\mathrm{e}}^{\mathrm{pool}}$; $\hat{\mathrm{C}}_{\mathrm{e}}^{\mathrm{df}}$ was used in **Fig. S3** and **S4** and **Main Fig.2**.

**Supplementary Note Table 3 | Effective Fig. S2 calibration denominators.** Values are minimum–median–maximum across the 20 groups in each band. Every group attempted 64 parameter settings with eight realizations per setting; a setting entered both response-surface fitting and residual pooling when it had at least two valid feature pairs. $N_e = \sum_i n_i$ is the number of pooled residual pairs and $D_e = \sum_i (n_i - 1)$ is the within-setting degrees of freedom. The quadratic design has 15 columns.

| Band | Estimator | Retained settings | $N_e$ | $D_e$ | Design rank |
|---|---|---|---|---|---|
| Alpha | FOOOF | 63–64–64 | 319–353.5–370 | 255–289.5–306 | 15–15–15 |
| Alpha | SPLIT | 25–33–40 | 67–90.5–111 | 42–59–73 | 15–15–15 |
| Beta | FOOOF | 64–64–64 | 393–413.5–427 | 329–349.5–363 | 15–15–15 |
| Beta | SPLIT | 37–42–50 | 110–123–152 | 72–81.5–102 | 15–15–15 |
| Low gamma | FOOOF | 63–64–64 | 418–431–449 | 354–367–385 | 15–15–15 |
| Low gamma | SPLIT | 39–44–55 | 108–127–152 | 68–84–97 | 15–15–15 |
| High gamma | FOOOF | 64–64–64 | 506–509–512 | 442–445–448 | 15–15–15 |
| High gamma | SPLIT | 64–64–64 | 371–391.5–414 | 307–327.5–350 | 15–15–15 |

All fitted quadratic design matrices had full column rank.

#### S2.4 Spearman correlation prediction and held-out evaluation

**Fig. S2c–f** reports Spearman rather than Pearson association, so the covariance matrix alone does not determine the displayed prediction.

We sampled one predicted conditional mean and one complete calibration residual independently, uniformly and with replacement. We added the selected vectors and repeated this procedure 50,000 times to predict the Spearman correlation between $P$ and $\chi$:

$$\tilde{\mathbf{z}}_b = \hat{\mathbf{m}}^{(2)}(\boldsymbol{\theta}_{H_b}) + \mathbf{e}_{E_b}, \qquad \rho_{\text{pred}} = \text{Spearman}(\{\tilde{P}_b\}, \{\tilde{\chi}_b\}) \tag{S15}$$

$b = 1, \ldots, 50{,}000$ indexed matched draws in which $H_b$ was sampled uniformly from estimator-specific held-out settings with at least one valid realization , $E_b$ selects one of the $N_e$ residual pairs,. Sampling each residual as a pair preserved the association between its peak-centre and exponent errors. Independent pairing makes the prediction distribution's covariance equal to $\hat{C}_m + \hat{C}_e$, with finite Monte

Carlo sampling error. It applies the pooled error distribution across input settings. The held-out pair was sampled uniformly from the valid realizations of that setting. The predicted pair combined the fitted second-order conditional mean at the same setting with one complete two-coordinate calibration residual sampled uniformly from the pooled residual distribution.

Using the same sampled setting indices makes both Spearman correlations setting-weighted. Sampling complete residual pairs preserves covariance between peak-frequency and exponent errors. For each estimator, a realization was successful when FOOOF or SPLIT yielded finite estimates of both periodic and aperiodic features. The observed successful fractions are reported below.

**Supplementary Note Table 4 | Successful held-out fitted pairs in Fig. S2.** Fractions are means across 20 groups per band; each group attempted 512 realizations.

| **Band** | **FOOOF fraction and range of valid counts** | **IRASA–SPLIT fraction and range of valid counts** |
|---|---|---|
| Alpha | 0.68330; 326–374 | 0.21445; 88–128 |
| Beta | 0.80586; 395–430 | 0.26963; 114–155 |
| Low gamma | 0.84258; 420–445 | 0.28770; 125–166 |
| High gamma | 0.99609; 507–512 | 0.76475; 376–410 |

The covariance ellipses in **Fig.S2c** used the first 4,096 predicted pairs and were translated to held-out means. In **Fig. S2f**, correction was the direct difference $\rho_{\mathrm{corrected}} = \rho_{\mathrm{held}} - \rho_{\mathrm{pred}}$. The difference quantifies residual association under the calibrated design.

Group summaries used 20 independently generated groups per band. One-sample two-sided Wilcoxon signed-rank tests compared the high-gamma group coefficients with zero; a paired test compared corrected FOOOF with IRASA–SPLIT in matching groups.

### S3 Estimator-specific influence functions

#### S3.1 Local feature derivatives with respect to log power

Equation (M5) in Methods asks which spectral frequencies make fitted peak centre and exponent move together, and Fig. S2b1,b2 displays the corresponding estimator sensitivities. This requires derivatives with respect to log-power bins rather than the generating parameters used in S2. Equations (S16)–(S19) derive the local rows $J_p$ and $J_{ap}$ while holding peak identity and active fitting constraints fixed.

The frequency-resolved analysis asks where a spectral perturbation changes an estimated feature. Let $L_j$ be log power at $f_j$ and let $F_{\boldsymbol{\eta}}(\mathbf{L})$ be estimator features on a fixed fitted branch, where $\boldsymbol{\eta}$ is a vector containing all estimated feature in the PSD fitting model. The formal definition is

$$J_{\boldsymbol{\eta}}(f_j) = \left.\frac{\partial F_{\boldsymbol{\eta}}(\mathbf{L})}{\partial L_j}\right|_{\mathbf{L}_0} \tag{S16}$$

For a spectral model $\ell_j(\boldsymbol{\eta}) = \ell(f_j\,;\boldsymbol{\eta})$ fitted by least squares, let $D_{jk} = \partial \ell_j / \partial \eta_k$. Differentiating its normal equations, while holding peak identities and active constraints fixed, gives a local sensitivity involving $D^{\top}D$ and residual-curvature terms. Neglecting those curvature terms gives the Gauss–Newton approximation

$$J_{\boldsymbol{\eta}} \simeq (D^{\top}D)^{\dagger}D^{\top}, \quad \delta\boldsymbol{\eta} \simeq J_{\boldsymbol{\eta}}\boldsymbol{\epsilon}. \tag{S17}$$

where $\epsilon_j$ denotes perturbations of original PSD model at frequency $f_j$. The dagger † is the Moore–Penrose pseudoinverse.

For knee-mode FOOOF, the fitted log spectrum is $b_{\mathrm{ap}} - \log_{10}(k + f^{\chi}) + \sum_l G_l(f)$. Derivatives of the aperiodic part are

$$\partial_{b_{\mathrm{ap}}}\ell = 1, \;\; \partial_k \ell = -\frac{1}{(k + f^{\chi})\ln 10}, \;\; \partial_{\chi}\ell = -\frac{f^{\chi}\ln f}{(k + f^{\chi})\ln 10}. \tag{S18}$$

Here $k$ is the fitted knee parameter. For $G = a\exp[-(f-c)^2/(2\sigma^2)] = ag$, the derivatives with respect to $a, c, \sigma$ are $g$, $ag(f-c)/\sigma^2$ and $ag(f-c)^2/\sigma^3$. The fitted bandwidth convention was converted consistently when forming these columns. All detected peaks were retained as nuisance terms in the local design matrix. The selected centre row and exponent row supply $J_p$ and $J_{ap}$.

SPLIT is treated differently from a joint parametric fit because its peak is estimated from original-minus-background log power while its exponent is estimated from the IRASA background.

Write $L_o$ and $L_f$ for the original and IRASA background log spectra. Peak fitting uses $L_o - L_f$, whereas exponent fitting uses $L_f$. If $J_{P,r}$ and $J_{\chi,f}$ denote their local rows, then

$$\delta\hat{P} = J_{P,r}(\delta L_o - \delta L_f), \quad \delta\hat{\chi} = J_{\chi,f}\delta L_f. \tag{S19}$$

where $\mathrm{P}$ is peak centre and $\chi$ is positive fitted aperiodic exponent. To obtain $J_{P,r}$ and $J_{\chi,f}$ in equation (S19), we fitted the oscillatory residual and the IRASA background as separate functions of their respective log-spectral inputs. Let $\mathrm{L_r} = \mathrm{L_o} - \mathrm{L_f}$ denote the original-minus-background log spectrum. The residual peak was represented by $\mathrm{G}(f_j\,; \mathrm{c}, \mathrm{h}, \sigma) = \mathrm{h}\exp[-(f_j - \mathrm{c})^2/(2\sigma^2)]$, with parameter vector $\eta_{\mathrm{P,r}} = (\mathrm{c}, \mathrm{h}, \sigma)^{\top}$. At the fitted reference peak, its local design matrix was $\mathrm{D_{P,r}} = \partial \mathrm{G}/\partial \eta_{\mathrm{P,r}}^{\top}$, whose columns were $\partial_{\mathrm{c}}\mathrm{G} = \mathrm{hg}(\mathrm{f}-\mathrm{c})/\sigma^2$, $\partial_{\mathrm{h}}\mathrm{G} = \mathrm{g}$ and $\partial_{\sigma}\mathrm{G} = \mathrm{hg}(\mathrm{f}-\mathrm{c})^2/\sigma^3$, where $\mathrm{g} = \exp[-(\mathrm{f}-\mathrm{c})^2/(2\sigma^2)]$. Applying the Gauss–Newton map in equation (S18), the peak-centre sensitivity was the centre row of the pseudoinverse, $\mathrm{J_{P,r}} = \mathrm{e_c^{\top}D_{P,r}^{\dagger}}$. The IRASA background was fitted independently as $\mathrm{L_f(f)} = \mathrm{b_f} - \chi\log_{10}\mathrm{f}$, with parameter vector $\eta_{\chi,\mathrm{f}} = (\mathrm{b_f}, \chi)^{\top}$ and linear design matrix $\mathrm{D}_{\chi,\mathrm{f}} = [1, -\log_{10}\mathrm{f}]$. Its exponent sensitivity was therefore $\mathrm{J}_{\chi,\mathrm{f}} = \mathrm{e}_{\chi}^{\top}\mathrm{D}_{\chi,\mathrm{f}}^{\dagger}$; this row is the exact ordinary-least-squares influence row for the fixed frequency grid. For $\mathrm{m}$ frequency bins, concatenating the two spectral inputs as $\mathrm{z} = (\mathrm{L_o^{\top}}, \mathrm{L_f^{\top}})^{\top} \in \mathbb{R}^{2\mathrm{m}}$ gives the joint first-order map

$$\mathrm{J_{SPLIT}} = \begin{pmatrix} \mathrm{J_{P,r}} & -\mathrm{J_{P,r}} \\ 0 & \mathrm{J}_{\chi,\mathrm{f}} \end{pmatrix}, \qquad \begin{pmatrix} \delta\hat{\mathrm{P}} \\ \delta\hat{\chi} \end{pmatrix} = \mathrm{J_{SPLIT}} \begin{pmatrix} \delta\mathrm{L_o} \\ \delta\mathrm{L_f} \end{pmatrix}. \tag{S20}$$

For SPLIT, the Gaussian peak and its peak-centre influence row $\mathbf{J_{P,r}}$ were fitted to the oscillatory residual L_r over the band-specific interval: 6–15 Hz for alpha, 12–35 Hz for beta, 25–70 Hz for low gamma and 50–170 Hz for high gamma. The aperiodic exponent and its straight-line influence row $\mathbf{J}_{\boldsymbol{\chi},\mathbf{f}}$ were obtained by ordinary-least-squares fitting of $\boldsymbol{L_f(f) = b_f - \chi\,\mathbf{log_{10}}\,f}$ over all bins of the common 0.2–200-Hz grid, with 0.2-Hz spacing.

#### S3.2 Frequency-resolved covariance contributions

**Fig. S2b3,b4** assigns signed covariance contributions to displayed frequencies after showing the normalized influence rows in panels **b1,b2**. Equation (S21) derives this decomposition from the full cross-frequency covariance $\Omega$ of the calibration spectra. Each displayed $K_j$ therefore includes covariance between bin $j$ and every other bin.

Across calibration spectra, sample centring gives the spectral covariance $\Omega$ at each frequency. Multiplying the two local feature perturbations yields

$$\mathrm{Cov}(\delta\hat{P},\delta\hat{\chi}) = J_p\Omega J_{ap}^{\mathsf{T}} = \sum_j K_j\,,\quad K_j = J_p(f_j)\sum_k \Omega_{jk}J_{ap}(f_k). \tag{S21}$$

Each $K_j$ contains the covariance of the error at $f_j$ with errors at every $f_k$. For SPLIT the same expression in block form is $J_{P,r}(\Omega_{of}-\Omega_{ff})J_{\chi,f}^{\mathsf{T}}$, where $\Omega_{of} = \mathrm{Cov}(L_o,L_f)$ and $\Omega_{ff} = \mathrm{Cov}(L_f)$.

Spectral covariance $\Omega$ was estimated from the representative high-gamma calibration group in Fig. S2b3,b4, which comprised 64 generating-parameter settings with 8 independent realizations per setting ($N = 512$ spectra). The spectra were centred across the entire pooled calibration group giving row matrices $X_o$ and $X_f$. Thus this diagnostic covariance contains both between-setting spectral variation and within-setting realization variation. For FOOOF, the covariance entering equation (S21) was

$$\hat{\Omega}_{oo} = \frac{X_o^{\mathsf{T}}X_o}{N-1}. \tag{S22}$$

For IRASA–SPLIT, the centred residual-log-spectrum matrix was $X_r = X_o - X_f$, and the effective residual–background cross-covariance was

$$\hat{\Omega}_{rf} = \frac{X_r^{\mathsf{T}}X_f}{N-1} = \hat{\Omega}_{of} - \hat{\Omega}_{ff} \tag{S23}$$

where $\hat{\Omega}_{of} = X_o^{\mathsf{T}}X_f/(N-1)$ and $\hat{\Omega}_{ff} = X_f^{\mathsf{T}}X_f/(N-1)$.

### S4 Two-dimensional stable-filter models and input synthesis

#### S4.1 Damped-oscillator and cubic-Hopf transfer functions

Equations (M11)–(M12) define both two-dimensional systems, their inputs and the parameter values used in the figures. For the linear system, the additional derivations begin by writing the linear part of equation (M11) in matrix form. Its unforced fixed point is the origin and

$$J_D = \begin{pmatrix} 0 & 1 \\ -\omega_0^2 & -2\gamma \end{pmatrix},\quad \det(\lambda I - J_D) = \lambda^2 + 2\gamma\lambda + \omega_0^2. \tag{S24}$$

Solving the characteristic polynomial gives $\lambda_\pm = -\gamma \pm \sqrt{\gamma^2-\omega_0^2}$. Thus $\gamma > \omega_0$ gives a stable node, $\gamma = \omega_0$ gives critical damping, and $0 < \gamma < \omega_0$ gives a stable complex pair. The transfer function is

$$H_D(\omega) = \frac{i\omega + 2\gamma}{\omega_0^2 - \omega^2 + 2i\gamma\omega},\quad \omega = 2\pi f. \tag{S25}$$

For the nonlinear cubic Hopf model, the Jacobian at the origin, its eigenvalue andthe corresponding linear transfer function are:

$$J_H = \begin{pmatrix} \mu & -\Omega \\ \Omega & \mu \end{pmatrix},\ \lambda_\pm = \mu \pm i\Omega,\ H_H(\omega) = \frac{i\omega - \mu}{(i\omega-\mu)^2 + \Omega^2}. \tag{S26}$$

#### S4.2 Bump-plus-aperiodic input generation

For stable-node regime in **Fig. S3b** and **Fig. S4b**, each input PSD is determined by $(\mathrm{c}, \mathrm{a}, \beta, \mathrm{b})$ through equation (M13). In each condition, two centred, unit-variance, orthogonal Gaussian vectors $u, v$ of length 64 defined $w = \rho_0 u + \sqrt{1-\rho_0^2}v$. Trial parameters were $(c, a, \beta, b) = (10 + 2.2u, 6 + 0.7u, -0.5 + 0.32w, 1 + 0.12w)$, clipped respectively to [5.5,18], [1,12], [-1.4,0.5] and [0.3,2]. In simulations, random phases $\phi \sim U(0,2\pi)$ is assigned, and inverse Fourier synthesis generated real inputs in time domain with the PSD shape in equation (M13). The zero-frequency component was zero, negative frequencies were conjugate symmetric, and each coloured trace was demeaned and normalized to RMS 0.02.

##### S4.3 Aperiodic-only input and condition selection

**Fig. S3a,b** and **Fig. S4a,b** use the bump-plus-aperiodic parameter ensembles defined in Equation (M13). **Fig. S3e,f** and **Fig. S4e,f** use aperiodic-only inputs. Equation (M14) gives the coloured-plus-white input form, whereas equation (S27) specifies its normalization and the text below gives the parameter ranges and condition-selection rule needed for reproduction.

To describe the normalization, define a unit-RMS coloured trace directly from its Fourier shape:

$$q_\beta(f_j) = \left[\frac{\max(f_j, 0.5)}{10}\right]^\beta, \quad v_n(\beta) = \mathrm{irFFT}\{\sqrt{q_\beta(f_j)}e^{i\phi_j}\}_n,$$
$$u_n(\beta) = \frac{v_n(\beta) - \bar{v}(\beta)}{\sqrt{N^{-1}\sum_m [v_m(\beta) - \bar{v}(\beta)]^2}}, \quad I_{i,n} = \sigma_I[u_n(\beta_i) + \nu\epsilon_{i,n}]. \tag{S27}$$

Here $\phi_j \sim U(0,2\pi)$, DC is set to zero before the inverse transform, and $\epsilon_{i,n} \sim \mathcal{N}(0,1)$ is independent of the coloured component. The input amplitude is $\sigma_I = 0.02$, and the independent white component has standard deviation $0.02\nu$. The sum is demeaned. This convention makes $u$ unit RMS and applies $\sigma_I$ once. It is algebraically the coloured-plus-white construction intended by equation (M14).

Within a condition, $\sigma_\beta$ and $\nu$ were fixed and $\beta_i$ followed a zero-centred Gaussian of scale $\sigma_\beta$, truncated to [−0.8,0.8]. The time-domain simulation used a Fourier grid extending to 2,000 Hz at 4,000-Hz sampling. The theoretical two-dimensional feature map normalized $(f/10)^\beta$ over 0.5–1,200 Hz on a 0.5-Hz grid before adding $\nu^2$. With a white floor, these panel-specific normalizations determine the relative coloured and white power as $\beta$ varies.

**In Fig. S4**, the theoretical prediction uses the transfer at the origin, the first-order parameter map, the three-point peak extractor and the Gamma-derived realization covariance. The simulation comparator uses the time-domain input in (S27), spline-based feature extraction and direct nonlinear cubic Heun trajectories. **Fig. S3** uses the exact linear transfer and the finite-window observation model specified in S6.4.

**Supplementary Note Table 5 | Additional two-dimensional sampling settings.** Model definitions and values already reported in Methods are not repeated. Rates expressed in hertz were multiplied by 2π in the differential equations.

| Setting | Oscillator | Cubic Hopf |
|---|---|---|
| Resonant damping candidates | $\gamma/\omega_0 \in [0.035, 0.55]$ | $\mu/(2\pi) \in [-8, -0.45]$ Hz |
| Other resonant candidates | $\sigma_\beta \in [0.05, 0.35]$; $\nu \in [0.20, 0.75]$ | Same |
| Candidate design | 5,000 Latin-hypercube conditions | Same |

The three Latin-hypercube coordinates were damping, $\sigma_\beta$ and $\nu$, using the model-specific damping coordinate in Table 4. The seed root was 20260809 with model-specific offsets. Acceptance required finite theoretical and observation-model correlations and theoretical $r < 0.6$. The first 1,000 accepted conditions, with 256 trials per condition, define the displayed analysis set.

### S5 State-space transfer and spectral feature sensitivity

#### S5.1 State linearization and input-parameter propagation

Equations (M18)–(M19) linearizes the extracted feature map with respect to input-generating parameters. **Fig. S3b,f, Fig. S4b,f** and **Main Fig. 2b,c** use both operations. Equations (S28)–(S29) derive the two operations separately so that a state Jacobian is not mistaken for a first-order parameter prediction.

Let a dynamical system be $\dot{\mathbf{x}} = F(\mathbf{x}, I(\mathrm{t}))$, where $I(t)$ denotes input, and $\mathbf{x}^*$ be a fixed point of this system at zero fluctuating input which satisfies $F(\mathbf{x}^*, 0) = 0$. Linearization gives $\dot{\delta \mathbf{x}} = J\delta\mathbf{x} + \mathbf{b}\, I(t)$ , where $J = \partial F / \partial \mathbf{x}|_{\mathbf{x}^*}$ and $\mathbf{b} = \partial F / \partial\, I(t)\, |_{\mathbf{x}^*}$. With output $y = \mathbf{v}^{\mathsf{T}}\delta\mathbf{x}$, Fourier transformation gives

$$\hat{y}(f) = \mathbf{v}^{\mathsf{T}}(2\pi i f I - J)^{-1}\mathbf{b}\ \ \hat{I}(f) = H(f)\hat{I}(f), \quad S_y(f) = |H(f)|^2 Q(f). \tag{S28}$$

Here $I$ within the matrix inverse is the identity matrix, $\hat{I}(f)$ is Fourier transform of input $I(t)$ and $Q(f)$ is input PSD. Stability requires negative real parts for every eigenvalue of $J$. For a linear system, this transfer is exact; for nonlinear systems it concerns local fluctuations around the selected fixed point.

Let $\boldsymbol{\psi}(\theta) = (P(\theta), A(\theta))^{\mathsf{T}}$ be extracted from the transfer-derived spectrum. A separate parameter expansion gives

$$\boldsymbol{\psi}(\theta) \simeq \boldsymbol{\psi}(\theta_0) + G(\theta - \theta_0), \quad C^{(1)}_{\mathrm{signal}} = G\Sigma_\theta G^{\mathsf{T}}. \tag{S29}$$

Rows of $G$ contain derivatives of peak frequency and signed slope. The covariance $\Sigma_\theta$ is calculated from the actual retained trial parameters. Adding calibration covariance and normalizing gives (M19), with the same algebra as (S15). If only $\beta$ varies, the signal covariance reduces to $C^{(1)}_{\mathrm{signal}} = \mathrm{Var}(\beta)\ \ \mathbf{g}_\beta \mathbf{g}_\beta^{\mathsf{T}}$. Without realization covariance or other varying parameters, this rank-one prediction gives correlation $+1$ or $-1$ when both derivatives are nonzero. Intermediate correlations can therefore depend strongly on the realization term.

#### S5.2 Peak-location and aperiodic-slope derivatives

Equations (M17)–(M18) define the peak and high-frequency slope extracted from each predicted output spectrum. The first-order predictions in Fig. S4b,f,  Fig. S5b,f and Main Fig. 2b,c require derivatives of those features. Equations (S30)–(S31) derive the implicit derivative of an interior peak and the linear log–log slope functional used in those calculations.

Suppose the smoothed log spectrum $\ell(f, \theta)$ has a unique interior maximum $P(\theta)$. Its defining conditions are $\ell_f(P, \theta) = 0$ and $\ell_{ff}(P, \theta) < 0$. Differentiating the first condition gives $\ell_{ff}|_{f=P}\, \partial_{\theta_k} P + \ell_{f\theta_k}|_{f=P} = 0$, hence

$$\partial_{\theta_k} P = -\frac{\ell_{f\theta_k}}{\ell_{ff}}|_{f=P}. \tag{S30}$$

For the aperiodic slope, put $x_j = \log_{10} f_j$ and $y_j = \log_{10} S_y(f_j)$. Minimizing $\sum_j (y_j - \alpha - Ax_j)^2$ over the chosen high-frequency bins gives

$$A = \frac{\sum_j (x_j - \overline{x})y_j}{\sum_j (x_j - \overline{x})^2} = \sum_j w_j\, y_j, \quad \frac{\partial A}{\partial \theta_k} = \sum_j w_j \frac{\partial y_j}{\partial \theta_k}. \tag{S31}$$

The slope band was 100–1,000 Hz in this study.

Numerical derivatives used the centred difference $[\boldsymbol{\psi}(\theta_0 + h_k e_k) - \boldsymbol{\psi}(\theta_0 - h_k e_k)]/(2h_k)$. The two-dimensional bump calculation used steps (0.02 Hz,0.02,0.002,0.002) at $(c, a, \beta, b) = (10{,}6, -0.5{,}1)$. Aperiodic-only derivatives used step $h_\beta = 0.002$ at $\beta = 0$.

### S6 Numerical evaluation and time-domain simulation

#### S6.1 Numerical routes and frequency-grid conventions

**Direct frequency-grid evaluation.** This route supplied the deterministic theory maps in **Fig. S3b,f** and **Fig. S4b,f**. Given an input spectrum, the transfer-derived output spectrum was calculated from equation (S28), with the finite-window Hann correction in equation (S41) where specified. For the two-dimensional theoretical maps, the unsmoothed log spectrum was evaluated on an evenly spaced 0.5-Hz grid from 0.5 to 1,200 Hz. The largest grid-bin log power in the prescribed peak band was refined by the three-point interpolation in the procedure in S6.5, and the signed high-frequency slope was calculated by equation (S31) over 100-1,000 Hz. Whole-brain transfer calculations in Fig. 2 used the same direct-grid principle but panel-specific grids specified in S8.

**Heun time-domain evaluation.** This route supplied the simulation results in Main **Fig. 2.** A sampled input sequence was propagated with the explicit Heun predictor-corrector update in equation (S32). Complete cubic or rectified simulations evaluated the nonlinear right-hand side at both Heun stages. For a fixed linear branch, the same update reduces to equation (S34) and can be evaluated by finite causal convolution. The resulting time series were converted to Welch spectra and analysed with the spline extractor in S6.5.

**Euler integration. Fig. S5** used direct rectified Euler integration, as specified in S8.1.

#### S6.2 Direct Heun integration and burn-in

For a differential equation with the sampled input held constant within each time step, explicit Heun integration used the following predictor and corrected update.

$$\mathbf{k}_1 = F(\mathbf{x}_n, I_n), \quad \tilde{\mathbf{x}}_{n+1} = \mathbf{x}_n + \Delta t \mathbf{k}_1,$$
$$\mathbf{k}_2 = F(\tilde{\mathbf{x}}_{n+1}, I_n), \quad \mathbf{x}_{n+1} = \mathbf{x}_n + \frac{\Delta t}{2}(\mathbf{k}_1 + \mathbf{k}_2). \tag{S32}$$

Direct network trajectories started from the fixed point of their own network structure. For an analysis input containing N samples, the final 2.048 s of the input were prepended as a warmup segment, and the corresponding  response was discarded. The integration step was $\Delta t = 0.25$ ms.

#### S6.3 Fixed-branch Heun recurrence and finite causal convolution

On a fixed active branch of the rectified network in Main Fig. 2b,c,f,g, perturbation dynamics are linear, with state Jacobian J and input vector b. Substitution of this linear drift into the Heun update gives the discrete recurrence below.

$$x_{n+1} = Mx_n + vI_n, \;\; M = I + \Delta t J + 1/2\,\Delta t^2 J^2, \;\; v = \Delta t b + 1/2\,\Delta t^2 J b. \tag{S33}$$

Here $M$ is the one-step state-update matrix and $v$ is the corresponding one-step input vector. Their calibration used the periodic linearized Heun response on the analysis-frequency grid, which is distinct from finite causal convolution. Held-out disconnection simulations in Main Fig. 2i instead integrated the complete rectified equations with equation (S32).

#### S6.4 Grid-based finite-window observation model

The finite-window observation model introduced below was used for both calibration and held-out evaluation of the linear oscillator, and to obtain the calibration covariance for the nonlinear Hopf analysis.

For each input-parameter vector, the exact linear transfer was evaluated on the frequency grid to obtain a deterministic output spectrum and feature pair. To approximate finite-record spectral variability, each frequency-bin power was multiplied by an independent random variable $V_j$

$$V_j \sim \text{Gamma}(k, k^{-1}), \quad k = \max\{3, 1.5(2T_a/4.096 - 1)\}. \quad (S34)$$

Thus, $\mathbb{E}[V_j] = 1$ and $Var(V_j) = 1/k$, with $k$ = 22.5 for the 32.768-s analysis and $k$ = 94.5 for the 131.072-s analysis. Perturbed log spectra were Gaussian-smoothed with standard deviation 1.95 bins on the 0.5-Hz grid before feature extraction. Peak frequency was then estimated by 3-point interpolation, and the signed aperiodic slope was fitted over 100–1,000 Hz (See Supplementary Notes S6.5). Feature perturbations from 1,024 spectral realizations for bump inputs, or 192 realizations for each aperiodic-only condition, defined the finite-window feature covariance. Zero-mean Gaussian feature errors with this covariance were added to the deterministic trial feature pairs. 3 independent observation realizations were used for calibration and 5 independent realizations were used for held-out evaluation. This procedure approximates finite-record feature variability without generating a time-domain ODE trajectory for each population-level trial.

#### S6.5 Peak and slope extraction from numerical outputs

**Frequency-grid spectra.** Transfer-derived and observation-model spectra used a three-point parabolic estimate around the largest grid-bin log power. The three adjacent log-power values and their frequency spacing determine the interpolated peak as follows:

$$\delta = \frac{l_- - l_+}{2(l_- - 2l_j + l_+)}, \quad P = f_j + \Delta f \ \ \text{clip}(\delta, -1, 1). \quad (S35)$$

where $l_j$ is the maximum of $log_{10}PSD$ in peak-search band and $f_j$ is the location of $l_j$ . If the magnitude of the underlying second difference in the denominator was at most 1 x $10^{-12}$, the interpolation offset was set to zero. The signed aperiodic slope was calculated by the OLS functional in equation (S31). Deterministic transfer maps were not smoothed unless a panel-specific finite-window observation or Hann-window operation was explicitly applied.

**Time-domain trajectories.** Direct dynamical outputs were analysed with 4.096-s Hann-windowed Welch segments, 50% overlap and constant detrending. Log power was Gaussian-smoothed with standard deviation 0.9765625 Hz, and a cubic spline was fitted to the smoothed log spectrum. The largest interior local maximum in the specified peak band defined the peak location $P$; the signed slope $A$ was fitted over 100-1,000 Hz.

**Validity masks.** Spline estimates were invalid when no peak was detected or when the selected peak lay within 0.5% of either search-band edge. Competing peaks differing in height by less than a factor of 1.20 were additionally excluded where specified. Whole-brain stable-node masks excluded competing-peak ambiguities; direct Hopf extraction excluded absent and boundary peaks without applying every competing-peak exclusion. At least 8 valid trials were required to calculate a correlation. Analysis-specific masks and stricter condition-selection rules are stated in S8.

**Supplementary Note Table 6 | Numerical procedures used for calibration and population evaluation.**

| Analysis | Calibration | Held-out evaluation |
|---|---|---|
| Two-dimensional linear oscillator (**Fig. S3**) | Gamma-derived Gaussian feature errors | Finite-window observation model |
| Cubic Hopf (**Fig. S4**) | Gamma-derived observation-model covariance | Direct Heun integration |
| Whole-brain stable node (**Fig. 2b,c**) | Full-network fixed-branch Heun | Finite causal fixed-branch Heun convolution |

| Analysis | Calibration | Held-out evaluation |
|---|---|---|
| Whole-brain stable complex **(Main Fig.2f,g**) | Condition-specific fixed-branch Heun | Finite causal fixed-branch Heun convolution |
| Whole-brain disconnection (**Main Fig. 2i,j**) | Linearized periodic Heun response | Direct Heun integration |
| 9-by-9 pathway supplement (**Fig.S5**) | Path-specific prediction calibration | Direct nonlinear Euler integration |

### S7 55-area network construction and local PG mode

#### S7.1 Anatomical connectivity and population parameters

Equation M19 defines the Li–Li–Wang marmoset network used throughout **Main Fig. 2** and **Fig. S5**. Each of 55 cortical areas contributes excitatory and inhibitory populations, so the simulated and linearized system has 110 states. Equations (S36) reconstruct the local and long-range weight blocks and the background drive.

The network uses the 55-area marmoset model, connectivity and hierarchy of Li, Li and Wang [3]. Let $F_{ij}$ denote an anatomical connection from source area $j$ to target area $i$, with zero diagonal. Let $h_i$ be hierarchical position normalized by its maximum. In state order $(\mathbf{r}_E, \mathbf{r}_I)$, the gain-scaled blocks of the matrix $W$ in (M22) are

$$\begin{gathered}(W_{EE})_{ij} = \beta_E(1+\eta_E h_i)(w_{EE}\delta_{ij} + \mu_{EE}F_{ij}),\\ (W_{EI})_{ij} = -\beta_E w_{EI}\delta_{ij},\\ (W_{IE})_{ij} = \beta_I(1+\eta_I h_i)(w_{IE}\delta_{ij} + \mu_{IE}F_{ij}),\\ (W_{II})_{ij} = -\beta_I w_{II}\delta_{ij}.\end{gathered} \tag{S36}$$

The Kronecker delta $\delta_{ij}$ selects local coupling. The subscripts on $W$ identify target and source population types. All long-range projections in this model originate from excitatory populations. The diagonal time-constant matrix $T$ has 55 entries $\tau_E$ followed by 55 entries $\tau_I$.

**Supplementary Note Table 7 | Whole-brain model constants.** Gain and coupling values use the source model's current and firing-rate convention; their products in (S38) are dimensionless. Time constants are converted together with frequency and integration units.

| Parameter | Value | Role |
|---|---|---|
| $\beta_E, \beta_I$ | 0.066, 0.351 | Excitatory and inhibitory population gains |
| $\tau_E, \tau_I$ | 20 ms, 10 ms | Population time constants |
| $w_{EE}, w_{EI}$ | 24.4, 19.7 | Local E-to-E and I-to-E strengths |
| $w_{IE}, w_{II}$ | 11.66038, 12.5 | Local E-to-I and I-to-I strengths |
| $\mu_{EE}, \mu_{IE}$ | 67.4, 49.81042 | Long-range E-to-E and E-to-I scales |
| $\eta_E, \eta_I$ | 0.685, 0.76 | Hierarchical modulation |

| Parameter | Value | Role |
|---|---|---|
| Baseline rates | 10 Hz E; 35 Hz I | Reference firing rates in all areas |

Scaling the local PG inhibitory-to-excitatory pathway multiplies its $W_{EI}$ entry by $S$ and constructs the corresponding baseline drive for that condition. In a disconnection comparison, the intact background is instead retained unchanged after removing long-range edges.

#### S7.2 Rectified fixed points and active sets

The network transfer in **Main Fig. 2** and **Fig. S5** is defined around a structure- and condition-specific fixed point of the rectified rate equations. Equation (S37) derives the solution on a candidate active branch and the accompanying sign checks needed before that branch can be used to construct a Jacobian.

For a network structure, solve $r^* = [Wr^* + b_0]_+$. Let $D$ be diagonal with entry one when the net drive $Wr^* + b_0$ is positive and zero when it is negative. Within a candidate branch the rectified equation is affine, so its fixed point satisfies

$$r^* = D(Wr^* + b_0), \quad r^* = (I - DW)^{-1} D b_0. \tag{S37}$$

#### S7.3 Full-network transfer and local PG node-to-focus diagnostic

Equations (M20)-(M21) use the full 110-state Jacobian and resolvent for the spectral predictions in **Main Fig. 2** and **Fig. S5**. **Main Fig. 2d** separately plots the eigenvalues of an isolated two-population PG block as the local inhibitory-to-excitatory multiplier changes. Equations (S38)–(S39) derive both objects and show why the discriminant crossing near $S = 2.08156$ is a stable node-to-focus transition rather than a Hopf bifurcation or a substitute for the full-network calculation.

Differentiating the drift $F(r, I) = T^{-1}\{-r + [Wr + b_0 + BI]_+\}$ at $(r^*, 0)$ gives the Jacobian and effective input vector used in M23–M24:

$$J = \left.\frac{\partial F}{\partial r}\right|_{r^*,0} = T^{-1}(-I + DW), \quad b_{\text{eff}} = \left.\frac{\partial F}{\partial I}\right|_{r^*,0} = T^{-1} D B. \tag{S38}$$

For output population selected by $v$, substitution into (S28) gives $H(f) = v^{\mathsf{T}}(2\pi i f I - J)^{-1} T^{-1} D B$. In millisecond-based code the frequency is $2\pi f/1000$ and the step is 0.25 ms; in second-based equations they are $2\pi f$ and 0.00025 s. These conventions are equivalent only when the Jacobian and time constants are converted consistently.

For an active E input, $T^{-1}DB = e_i/\tau_E$. Some continuous spectral-feature routines omit this constant input factor because it changes power scale but not peak frequency or log slope. The time-domain equations retain it. If the input population is inactive, its local forcing derivative vanishes; one cannot replace $DB$ by $B$ indiscriminately.

The local PG block diagnoses the local real-to-complex transition. Writing its characteristic polynomial as $\lambda^2 - (\mathrm{tr}J_{PG})\lambda + \det J_{PG} = 0$ and applying the quadratic formula gives

$$\lambda_\pm(S) = \frac{\mathrm{tr}J_{PG}}{2} \pm \frac{1}{2}\sqrt{(\mathrm{tr}J_{PG})^2 - 4\det J_{PG}}. \tag{S39}$$

A negative discriminant gives a complex pair, with real part $\mathrm{tr}J_{PG}/2$. Hence the discriminant crossing is a node-to-focus transition. It is not a Hopf bifurcation unless the real part also reaches zero with a nonzero imaginary part. The complex onset used in **Main Fig. 2d** was near $S = 2.08156$.

For any real frequency $f$, $(2\pi i f I - J)^{-1} = \mathrm{adj}(2\pi i f I - J)/\det(2\pi i f I - J)$. If the coalesced eigenvalue remains in the open left half-plane, the denominator is nonzero on the imaginary axis. Matrix entries of the resolvent, the transfer $H(f)$ and the transfer-derived PSD therefore remain smooth functions of $S$ through the stable coalescence. A discontinuity in extracted $P$ would instead indicate peak switching, a search boundary or another feature-extraction branch change. The spectral predictions therefore used the full resolvent, rather than differentiating an individually labelled eigenvalue. The local block is not the 110-state Jacobian: full-network stability was assessed separately, and the full-network transfer was used for network prediction.

### S8 Whole-brain prediction and structural disconnection

#### S8.1 Resting stable-node input–output map

**Main Fig. 2b** asks whether a first-order theory predicts coupling across 1,000 independent 64-trial input ensembles in the resting network, while panel c imposes zero realized input association and **Fig. S5** repeats the pathway analysis for nine inputs and nine readouts. Equation (S40) records the coordinate map about $\theta_0$, after which the calibration design and realization covariance used by these panels are specified.

The stable-node experiment asks how a fixed resting network transmits input-feature association. Connectivity, background and the V1 input to PG output pathway were fixed. Trials varied the parameters of the bump-plus-aperiodic spectrum in (M16). Prediction used the first-order parameter covariance (S29) about $\theta_0 = (10,6,-0.5,1)$, plus calibration-derived realization covariance. This separates input association from changes in the network itself.

**Input construction.** Input coordinates $\theta$ in **Main Fig. 2b** were constructed by adding a linearly transfromed perturbation term $z$ on the baseline parameter $\theta_0$:

$$\theta = \theta_0 + D_\theta z\,, \quad D_\theta = \mathrm{diag}(3,4,0.8,0.7). \tag{S40}$$

The bounds of $\theta$ were $(5.1,1,-1.8,0.15) < \theta < (19.5,15,1,3)$. The main 1,000-condition test in **Main Fig. 2b** set $z = (0.2u, 0, 0.2w, 0)$, where $u,w$ are calculated as in S4.2. Input PSDs $Q(f;\theta)$ were constructed by equation (M16). Input time series were then constructed by $x_k(t,\theta) = a_0 \mathrm{irFFT}\{\sqrt{Q(f;\theta)}\, e^{i\phi_k}\}$with DC removed, where $k$ denotes different realizations under a trial, $\phi_k \sim U(0,2\pi)$ is a random phase and $a_0 = 0.5/[\mathrm{SD}_N(x_k(t,\theta_0)) + 10^{-15}]$ is a common multiplier.

The separate zero-input-association control in **Main Fig. 2c** used 100 independently constructed 64-trial ensembles. Iterative projection adjusted the colour coordinate until the correlation of extracted input features had magnitude below $10^{-12}$.

**Transfer-derived Jacobian.** The transfer-derived Jacobian of the feature map $G = \frac{\partial(P,A)}{\partial \mathbf{z}}$ was used in theoretical prediction in **Main Fig. 2**b,**c**. To obtain $G$, output PSD was calculated by the transfer function in equation (M21) and input PSD on the specified frequency grid (The frequency grid contained 180 geometrically spaced points from 0.5 to 40 Hz and 220 from 40.1 to 1,200 Hz), and periodic feature $P$ and aperiodic feature $A$ were then extracted. $G$ was then calculated with step 0.002 in each coordinate of $\mathbf{z}$ using the central difference method.

**Calibration.** Calibration used 3 realizations of 14 independent 64-trial ensembles at standardized scale 0.2. Each ensemble contained 64 trial-level parameter settings in the coordinate order $z_1$=bump centre, $z_2$=bump height, $z_3$=aperiodic colour and $z_4$=aperiodic amplitude. The parameter distribution for each ensemble is listed in Supplementary Note Table 7. Standardized coordinates were then converted to physical parameters using equation (S40) and then used to construct the bump-plus-aperiodic input spectrum. 3 independent Fourier-phase realizations were generated for every setting. Within each setting, the resulting output feature pairs were centred across valid realizations, and their complete two-dimensional residual outer products were pooled over the 14 ensembles using the total within-setting degrees of freedom as the denominator. This single-realization covariance therefore captures variability from random Fourier phases, finite records, Welch estimation and feature extraction while the generating parameters are fixed.

For the main V1-to-PG stable-node map (**Main Fig. 2b**), the pooled calibration covariance from these 14 ensembles was added to the first-order signal covariance for each of the 1,000 independent 64-trial evaluation ensembles. The zero-association control (**Main Fig. 2c**) instead estimated a separate condition-specific covariance from its own three calibration realizations. The archived nine-by-nine pathway analysis (**Fig. S5**) used the same 14 design families but estimated covariance separately for each input-output pathway and design.

**Supplementary Note Table 8 | Whole-brain stable-node regime calibration design.**

| No. | Design | Standardized-coordinate construction |
|---|---|---|
| 1 | Independent Gaussian | $z_i = 0.2\xi_i, \quad \xi_i \sim \mathcal{N}(0,1)$ |
| 2 | Rank one | $z_i = 0.2t_i v_i, \quad v = (1,0.6,-0.8,0.4)^T, \quad t_i \sim \mathcal{N}(0,1)$ |
| 3 | Uniform | $z_i = 0.2\xi_i, \quad \xi_i \sim U(-\sqrt{3},\sqrt{3})$ |
| 4 | Fixed total | $z_i = 0.2(\xi_1,\xi_2,\xi_3,-\xi_2)^T, \quad \xi_i \sim \mathcal{N}(0,1)$ |
| 5 | Standardized lognormal | $x_i = Lognormal(0,0.55^2), \quad \xi_i = (x_i - \bar{x})/SD(x_i), \quad z_i = 0.2\xi_i$ |
| 6 | One-factor mixture | $k_i = i \bmod 4, \quad z_{i,k_i} = 0.2\xi_i, \quad \xi_i \sim \mathcal{N}(0,1), \quad z_{i,j} = 0\ for\ j \neq k_i$ ($i$ is trial number) |
| 7 | Second Gaussian realization (same as ensemble 1, with a different random seed) | $z_i = 0.2\xi_i, \quad \xi_i \sim \mathcal{N}(0,1)$ |
| 8 | Correlated Gaussian $\rho = -0.9$ | $z_i = 0.2\xi_i, \quad \xi_i \sim \mathcal{N}(0,1), \quad Corr(\xi_1,\xi_3) = -0.9$ |
| 9 | Correlated Gaussian $\rho = -0.6$ | $z_i = 0.2\xi_i, \quad \xi_i \sim \mathcal{N}(0,1), \quad Corr(\xi_1,\xi_3) = -0.6$ |
| 10 | Correlated Gaussian $\rho = -0.3$ | $z_i = 0.2\xi_i, \quad \xi_i \sim \mathcal{N}(0,1), \quad Corr(\xi_1,\xi_3) = -0.3$ |
| 11 | Correlated Gaussian $\rho = 0$ | $z_i = 0.2\xi_i, \quad \xi_i \sim \mathcal{N}(0,1), \quad Corr(\xi_1,\xi_3) = -0$ |
| 12 | Correlated Gaussian $\rho = 0.3$ | $z_i = 0.2\xi_i, \quad \xi_i \sim \mathcal{N}(0,1), \quad Corr(\xi_1,\xi_3) = 0.3$ |
| 13 | Correlated Gaussian $\rho = 0.6$ | $z_i = 0.2\xi_i, \quad \xi_i \sim \mathcal{N}(0,1), \quad Corr(\xi_1,\xi_3) = -0.6$ |
| 14 | Correlated Gaussian $\rho = 0.9$ | $z_i = 0.2\xi_i, \quad \xi_i \sim \mathcal{N}(0,1), \quad Corr(\xi_1,\xi_3) = 0.9$ |

**Full-network simulation.** Full-network fixed-branch Heun convolution produced 32.768-s analysis records after 2.048-s burn-in. The main map used one evaluation realization per condition. The zero-association control used three calibration and five held-out realizations per condition and averaged the five held-out correlations, without dividing the single-realization covariance by five. Peak extraction used 5–20 Hz. The nine-by-nine pathway extension varied the input and output areas over V1, V2, V4, MST, LIP, PG, A23a, A6DR and A6Va; its population simulations used direct Euler integration, as distinguished in Table 5.

#### S8.2 Stable-complex complete feature-map prediction

**Main Fig. 2d–g** examine a resonant stable network driven by aperiodic-only inputs. For every retained condition, the fixed point and full transfer are recomputed, and each of 256 trial spectra is propagated through the Hann-corrected transfer before its peak and slope are extracted. Equation (S41) gives the Hann-window correction and equation (S42) combines the covariance of those complete trial-wise feature pairs with condition-specific realization covariance.

**Hann-window spectral spreading.** Methods M25 predicts the stable-complex spectra shown in **Main Fig. 2e–g** from a finite Hann-windowed record. Equation (S41) expresses the corresponding spreading as a convolution of the transfer-derived even spectrum with the squared periodic-Hann response on the signed FFT grid. This corrects the deterministic mean spectrum; realization covariance is added separately.

A finite spectral window spreads power across nearby frequencies. If $w_n$ is a length-$L$ analysis window, the expected windowed periodogram of a stationary signal is proportional to the convolution of its two-sided PSD with $|W(f)|^2$, where $W$ is the Fourier transform of $w$. Welch averaging reduces variability but retains this window-induced mean. On a discrete frequency grid the normalized representation is

$$K_{jk}^{\mathrm{win}} = \frac{|W(f_j - f_k)|^2}{\sum_l |W(f_j - f_l)|^2}, \quad \bar{S}_y(f_j) = \sum_k K_{jk}^{\mathrm{win}} \, |H(f_k)|^2 Q(f_k). \tag{S41}$$

For the periodic Hann window, $w_n = [1 - \cos(2\pi n/L)]/2$. The whole-brain stable-complex calculation used $L = 16{,}384$ at 4,000 Hz, with a signed FFT grid of length $4L = 65{,}536$. Transfer magnitude and input shape were evaluated at absolute frequency, forming an even two-sided spectrum. Circular FFT convolution was performed on the complete signed grid. Every fourth positive-frequency bin then recovered the Welch grid, before restricting to the analysis range.

**Input sampling criteria.** An input condition is determined by PG inhibitory-to-excitatory coupling multiplier $S$(range: 3.2662-6.6938), input colour spread $\sigma_\beta$ (range: 0.05022-0.41095) and white-noise ratio $\nu$ (range: 0.07207-0.49997). Latin-hypercube design sampled $(S, \sigma_\beta, \nu)$ for each condition with additional enriched candidates. The candidate ensemble comprised a 2,200-condition Latin hypercube and 3,000 enriched candidates. Table 7 gives the two candidate domains; design seeds were 20260820 and 20260821. Candidates were examined in a fixed order and required finite prediction and simulation statistics, at least 80 common valid trials and theoretical $r < 0.6$. The first 1,000 accepted conditions define the window-corrected theoretical and Heun simulation analyses. The retained ensemble contained 1,000 conditions with 256 trials each.

**Supplementary Note Table 9 | Stable-complex candidate domains.** The retained ranges are reported in Methods and are not repeated here. These domains describe condition-level parameters; trial-level $\beta$ is sampled separately within each condition.

| Parameter | Base candidates | Enriched candidates |
|---|---|---|
| $S$ | 2.081563–6.694 | 4.5–6.694 |

| Parameter | Base candidates | Enriched candidates |
|---|---|---|
| $\sigma_\beta$ | 0.05–0.6 | 0.05–0.25 |
| $\nu$ | 0–0.5 | 0.3–0.5 |

The full fixed point was active in all 110 coordinates in the retained baseline conditions. This statement concerns equilibria, not a proof that every noisy trajectory remained active. Every retained full-network Jacobian was stable. The coloured input had RMS 0.02 Hz and white-noise standard deviation $0.02\nu$ Hz.

3 calibration realizations estimated the condition-specific, degrees-of-freedom-corrected single-realization covariance in (S42). 5 independent held-out realizations regenerated phases and white samples using the same trial parameters. Their 131.072-s records followed 2.048-s burn-in, using fixed-branch Heun convolution at 0.25 ms. The peak band was 5–100 Hz. Candidate construction retained conditions with at least 80 common valid trials. After the 1,000-condition ensemble was frozen, a condition-level correlation was defined only when at least eight trials remained under the common validity maskt. Simulation correlation was calculated across trials separately in each held-out realization, then averaged over the five values. It was not the correlation of averaged PSDs or pooled realization pairs.

**First-order expansion theory.** For each of the 1,000 frozen network conditions, we recomputed the fixed point, active set, Jacobian and full-network transfer function. Each trial input spectrum was propagated through this transfer and the finite Hann-window kernel. We extracted peak frequency *P* and signed log–log slope *A*, and defined the aperiodic exponent as *χ*=-*A*. The displayed theory then expanded the two-feature map $\mathbf{m}(\beta) = (P(\beta), \chi(\beta))^T$ to first order in the trial-level input-colour parameter β about $\beta_0 = 0$.

$$m(\beta) \approx m(0) + g\beta, g = (\partial m/\partial\beta)|_{\beta=0}, C_{signal}^{(1)} = gVar(\beta)g^{\mathrm{T}} \tag{S42}$$

We estimated g by centred finite differences with stepsize $h$ = 0.002; fixed checks at $h$ = 0.004 and 0.001 verified convergence.

**Covarian**ce prediction. For each condition, the network fixed point, Jacobian and transfer were recomputed. Each trial's input spectrum was passed through the full transfer and Hann-window kernel in (S41). Extracting a pair $m_i = (P_i, A_i)^{\mathrm{T}}$ from each predicted spectrum gave

$$\hat{C}_m = \frac{1}{n-1}\sum_{i=1}^{n}(m_i - \overline{m})(m_i - \overline{m})^{\mathrm{T}}, \quad C = \hat{C}_m + \hat{C}_e^{\mathrm{df}}, \quad r_{\mathrm{pred}} = \frac{C_{PA}}{\sqrt{C_{PP}C_{AA}}}. \tag{S43}$$

This reproduces the complete trial-wise feature-map calculation in (M25)–(M26). It linearizes network dynamics about the fixed point but does not expand the feature map to first order in $\beta$.

#### S8.3 Whole-brain structural disconnection

Equations (M27)–(M28) compare intact and deleted network structures in **Main Fig. 2h1,h2,i,j.** Deletion removes the selected areas' incoming and outgoing long-range edges but retains their local circuits and all 110 state variables. Equations (S44) define the signed change between two structure-specific complete feature-map predictions, the realization term appropriate to within-trial PSD averaging, the paired input covariance and the direct-output indicator used in the displayed comparisons.

**Input Construction.** Inputs were delivered to A6Va, using (M16), the centre and coordinate scaling in (S40), and the whole-brain clipping bounds in S8.1. Standardized perturbations followed $z \sim \mathcal{N}(0, 0.3^2\Sigma_d)$. In $(c, a, \beta, b)$ order it was approximately

$$\Sigma_d = \begin{pmatrix} 0.01225258 & 0.00018177 & -0.04698533 & -0.00019086 \\ 0.00018177 & 0.01003743 & -0.00382986 & -0.00001556 \\ -0.04698533 & -0.00382986 & 1 & 0.00402134 \\ -0.00019086 & -0.00001556 & 0.00402134 & 0.01003895 \end{pmatrix}.$$

Its condition number was 100 before sampling and clipping, although the finite-sample covariance can differ. Each 512-trial ensemble was generated by matched random sampling, with common trial vectors and phase seeds pairing the intact and disconnected responses.

**Supplementary Note Table 10 | Disconnection evaluation designs.** Realization counts are per trial. Different windows and averaging counts define different observation precision.

| Design | Theory–simulation scatter in Main Fig.2i | Direct versus non-direct node-count comparison in Main Fig. 2j |
|---|---|---|
| Patterns | 82 selected patterns | 60 patterns, 12 for each deletion count 1–5 |
| Trials per pattern | 128 | 512 |
| Calibration realizations | 3 | 2 |
| Held-out realizations $H_r$ | 5 | 2 |
| Analysis window | 32.768 s | 16.384 s |
| Integration | Direct rectified Heun, 0.25 ms | Same |
| Burn-in | 2.048 s | Same |
| Readout | V2 | Eligible non-deleted E outputs |

**Node deletion.** The 82-pattern A6Va-to-V2 scatter in **Main Fig. 2i** comprised 48 one-area, 19 three-area, 12 four-area and 3 five-area patterns. Selection retained fixed-point-active input and output. For the node-count comparison, the candidate pool was balanced by V2's direct connectivity and supplemented by stable patterns with active A6Va and V2. 12 patterns were sampled without replacement for each count, with seed 20260930.

**Calibration and held-out simulation.** Calibration used the fixed-branch Heun response in S6.3. Held-out simulations started at each structure's fixed point and integrated the complete rectified equations, as stated in S6.2. Within each trial, held-out PSDs were averaged before extracting one $(P, A)$ pair. Correlations across trials were computed separately for intact and disconnected structures, then subtracted.

Validity was applied in a fixed hierarchy. First, a structure–output pair was eligible only if the input and output area were not deleted and were active at the relevant modified fixed point. Second, each trial is valid only when the aperiodic feature $A$ and periodic feature $P$ are detected and finite in both intact network and node-deleted network.

**Definition of Direct and non-direct outputs.** An eligible output was neither deleted nor inactive at the modified fixed point. Using the intact directed anatomical matrix, direct status was

$$d_o(D_s) = \mathbf{1}\{\exists j \in D_s : F_{oj} > 0\}. \tag{S44}$$

Thus an output was direct if it received an edge from any deleted area. For each pattern and source, the reported class statistic was the mean of $|\Delta r_o|$ over eligible outputs in that class.

For each deletion pattern, $|\Delta r \quad |$ was averaged across eligible outputs separately within the direct and non-direct classes. Deletion patterns were the paired statistical units. Patterns with an empty direct or non-direct class were excluded from that comparison, and the retained number of paired patterns was reported separately for each deletion count. The resulting output- and network-level denominators are given below. Counts are identical for theory and simulation because their output-level table used the common eligibility structure.

**Supplementary Note Table 11 | Effective output and paired-pattern denominators for the node-count comparison.**

| Deleted areas | Direct outputs; patterns | Non-direct outputs; patterns | Paired network means |
|---|---|---|---|
| 1 | 413; 12 | 231; 12 | 12 |
| 2 | 502; 12 | 121; 12 | 12 |
| 3 | 532; 12 | 74; 11 | 11 |
| 4 | 517; 12 | 45; 10 | 10 |
| 5 | 504; 12 | 52; 12 | 12 |
